# High-resolution simulations of chromatin folding at genomic rearrangements in malignant B-cells provide mechanistic insights on proto-oncogene deregulation

**DOI:** 10.1101/2021.03.12.434963

**Authors:** Daniel Rico, Daniel Kent, Nefeli Karataraki, Aneta Mikulasova, Rolando Berlinguer-Palmini, Brian A. Walker, Biola M. Javierre, Lisa J. Russell, Chris A. Brackley

**Author notes:** Senior authors. Address correspondence to, or.

## Abstract

Genomic rearrangements are known to result in proto-oncogene deregulation in many cancers, but the link to genome 3D structure remains poorly understood. Here we used the highly-predictive heteromorphic polymer (HiP-HoP) model to predict chromatin conformations at the proto-oncogene *CCND1* in healthy and malignant B-cells. After confirming that the model gives good predictions of Hi-C data for the non-malignant human B-cell derived cell line GM12878, we generated predictions for two cancer cell lines, U266 and Z-138. These possess genome rearrangements involving *CCND1* and the immunoglobulin heavy chain locus (*IGH*), which we mapped using targeted genome sequencing. Our simulations showed that a rearrangement in U266 cells where a single *IGH* super-enhancer is inserted next to *CCND1* leaves the local topologically associated domain (TAD) structure intact. We also observed extensive changes in enhancer-promoter interactions within the TAD, suggesting that it is the downstream chromatin remodelling which gives rise to the oncogene activation, rather than the presence of the inserted super-enhancer DNA sequence *per se*. Simulations of the *IGH*-*CCND1* reciprocal translocation in Z-138 cells revealed that an oncogenic fusion TAD is created, encompassing *CCND1* and the *IGH* super-enhancers. We predicted how the structure and expression of *CCND1* changes in these different cell lines, validating this using qPCR and fluorescence *in situ* hybridization microscopy. Our work demonstrates the power of polymer simulations to predict differences in chromatin interactions and gene expression for different translocation break-points.

---

Chromatin structure, nuclear organisation and the epigenome are intimately linked to gene regulation and cell function, and are tightly controlled during differentiation. Genome rearrangements leading to structural variants (including deletions, insertions, duplications and translocations) can disturb these, leading to gene mis-regulation and cancer (Norton and Phillips-Cremins, 2017). An important example occurs during B-cell differentiation, where programmed breakage and recombination of the genome takes place in order to generate the broad heterogeneity of immunoglobulins (Igs) required for immune system function (Jung *et al*., 2006). Errors in this process can lead to repositioning of Ig regulatory elements which then drives proto-oncogene activation (Kontomanolis *et al*., 2020). The diversity in translocations and the difficulties in accessing and handling patient samples means that detection, accurate mapping, and characterisation of their functional consequences are inherently problematic.

Advances in molecular probes of genomic and epigenomic structure, such as ChIP-seq and chromosome-conformation-capture methods like Hi-C, have transformed our understanding of the regulatory link between structure and function. For example, Hi-C, which probes chromosome interactions genomewide, has revealed that the genome can be partitioned into topologically associated domains (TADs) (Dixon *et al*., 2012).

These are contiguous chromosome regions which show enriched self-interactions, and are thought to be associated with cis-regulatory mechanisms; they are often bounded by binding sites for the CCCTC-binding factor CTCF (Rao *et al*., 2014). ChIP-seq profiling of histone modifications and protein binding has identified super-enhancers, clusters of enhancer elements thought to drive expression patterns responsible for cell identity (Hnisz *et al*., 2013). Super-enhancers have been found to preferentially sit within TADs which are highly insulated from their surroundings (Gong *et al*., 2018), and have been identified as drivers of oncogene expression in many tumours (Thandapani, 2019; Mikulasova *et al*., 2020b).

A full epigenetic and three-dimensional (3D) structural characterisation of genome rearrangements is crucial for our understanding of proto-oncogene activation. However, the small size of patient samples and their inherent variability means that routinely applying methods like Hi-C to primary cancer samples remains challenging, despite recent progress in reducing the amount of material required (Díaz *et al*., 2018). This is why much work to date has focused on the biogenesis of chromosome translocations (Engreitz *et al*., 2012; Roukos and Misteli, 2014), but comparatively less attention has been paid to the structural *consequences* of translocations and other genome rearrangements (Bianco *et al*., 2018; Akdemir *et al*., 2020).

In this work, our aim was to show how computer simulations can help us understand the effects of common genome rearrangements in B-cell malignancies. Specifically, we used our “highly-predictive heteromorphic polymer” (HiP-HoP) model (Buckle *et al*., 2018; Brackey *et al*., 2020) to study genomic rearrangements involving the Ig heavy chain locus (*IGH*) (Watson and Breden, 2012) and the *CCND1* proto-oncogene encoding the cell cycle protein cyclin D1. *IGH*-*CCND1* is a common translocation observed in mantle cell lymphoma (MCL) and multiple myeloma (MM); when translocated together the *IGH* superenhancers drive *CCND1* overexpression. The goal was to use the model to shed light on how such rearrangements lead to changes in the 3D structure of gene loci which in turn leads to deregulation. If these simulations allow us to infer the pathway through which the *CCND1* proto-oncogene becomes activated, they could be used to generate testable hypotheses to direct new experiments suggesting targets for therapeutic intervention. In the longer term, this points to a future role for the method in a clinical setting, *e*.*g*., in personalised medicine, where structural predictions based on available or easily obtainable data could be extremely valuable.

## Results

Figure 1 shows a cartoon of the *IGH* and *CCND1* loci in healthy cells, as well as two different rearrangements which are studied here. Specifically we consider an MM cell line, U266, where a single *IGH* super-enhancer is inserted into the *CCND1*-TAD, and an MCL cell line, Z-138, possessing a translocation which repositions *CCND1* next to the *IGH* joining region (Watson and Breden, 2012). In previous work (Mikulasova *et al*., 2020b) we analysed the changes in patterns of histone modifications associated with super-enhancer translocation events, finding that relocation of the *IGH* super-enhancer results in local chromatin remodelling and the emergence of new patterns of histone modifications including H3K4me3 broad domains (Park *et al*., 2020). Here we study how these epigenetic changes alter 3D chromosome structure.

**Figure 1.**
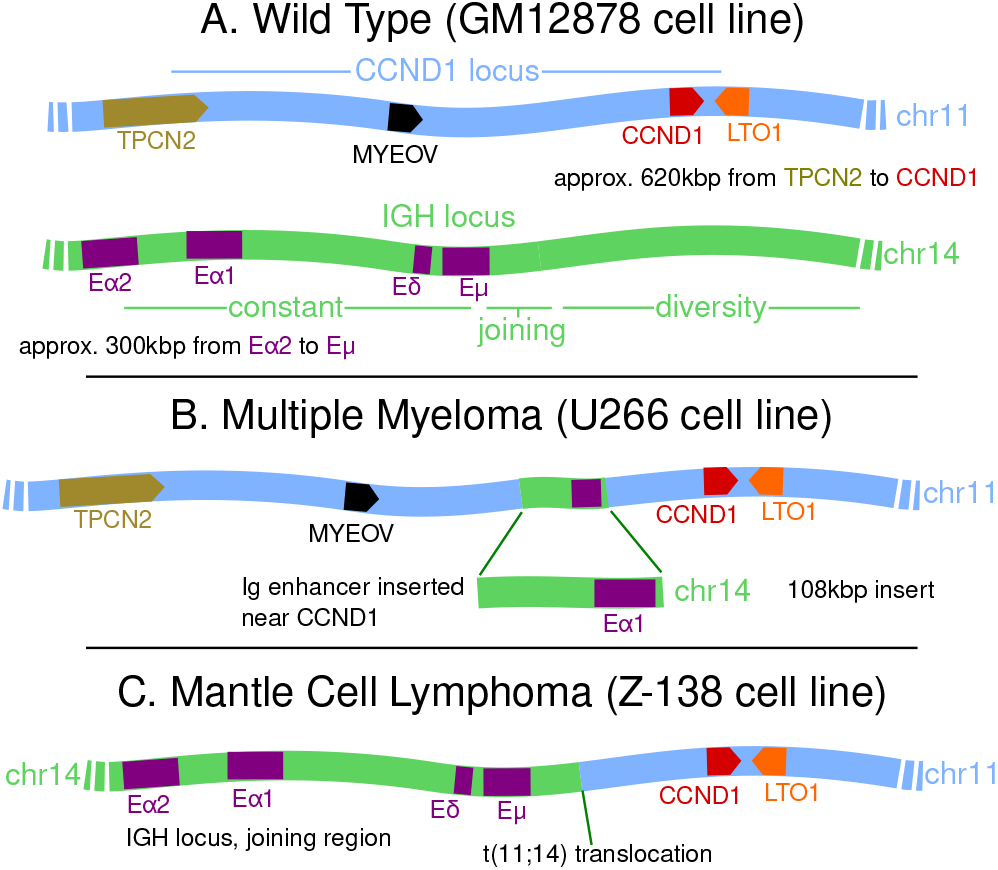
Cartoons showing the layout of the *CCND1* and *IGH* loci. **A**. Healthy cells. For *CCND1* we show the approximate positions and orientations of four genes within the TAD (*TPCN2* lies just outside the TAD which encloses the other three). For *IGH* we show four previously identified enhancer elements within the constant region. **B**. A rearrangement which has been observed in multiple myeloma (and in the U266 cell line) where an *IGH* insert bringing an enhancer element into the vicinity of *CCND1* is shown. **C**. A t(11;14) translocation often found in mantle cell lymphoma (and in the Z-138 cell line) which brings *CCND1* next to the entire *IGH* constant region is shown.

HiP-HoP uses data on DNA accessibility, histone modifications, and protein binding to generate an ensemble of 3D chromosome structures. We previously showed that the method gives good predictions of both population level data (e.g., Hi-C or 4C) and single cell measurements [e.g., from fluorescence microscopy; Buckle et al. (2018)]. Importantly, HiP-HoP does not use Hi-C (or any other 3D genome data) as an input [unlike other popular models which use fitting-based methods; Serra et al.(2015)]. This means that we can make predictions about the 3D structure of cell types or tissues where Hi-C data is not available.

### HiP-HoP predicts the domain structure at CCND1 in a B-cell derived cell line

We previously showed that in malignant B-cells, genome rearrangements involving CCND1 lead to local changes in histone modifications (Mikulasova et al., 2020b). Here we hypothesised that this in turn leads to changes in 3D structure and chromatin interactions which drive *CCND1* expression. To understand these malignant structural changes, it is first necessary to study the healthy case. To this end we considered the human B-cell derived lymphoblastoid cell line GM12878, adapting our HiP-HoP method to study chromatin structure around *CCND1*. This is an ideal cell line due to the abundance of publicly available data.

In the model, a 3 Mbp chromosome region was represented by a chain of beads [a common approach in polymer physics; Brackey *et al*. (2020)]. The region was chosen because it is large enough to encompass seven TADs surrounding *CCND1* such that the local chromatin context is captured, but small enough to give feasible simulation run times. The model combines three mechanisms which drive locus structure: (i) diffusing spheres representing complexes of chromatin binding proteins which can bind to multiple points at the same time to form molecular bridges (*e*.*g*., between promoters and enhancers) (Brackley *et al*., 2013, 2016b; Brackley, 2020); (ii) a heteromorphic polymer structure, where different sections of the bead chain have different properties (thickness/compaction and flexibility); and (iii) the loop extrusion mechanism (Fudenberg *et al*., 2016; Sanborn *et al*., 2015). Loop extrusion is a molecular mechanism where chromatin is pushed onto loops by factors [the cohesin complex; Davidson *et al*. (2019)], which are halted by CTCF proteins bound in a convergent orientation (Rao *et al*., 2014).

A way to identify different types of chromatin is via highthroughput analysis of histone modification data and hidden Markov modelling (HMM) (Ernst *et al*., 2011; Carrillo-de SantaPau *et al*., 2017); this approach classifies chromosome regions into a number of states (*e*.*g*., promoter/enhancer associated, polycomb repressed, and heterochromatin states, see Supplemental Table S1). In the model we had two different types of chromatin structure (the heteromorphic polymer): more open regions (thinner, more flexible polymer) and more compact regions (thicker, “crumpled” polymer, see Supplemental Fig. S1 for a schematic). We used the chromatin states to identify these regions: H3K27ac associated states have the more open structure (Risca *et al*., 2017). The model included three different species of bridge forming protein: a general “active binder”, and two types of repressive binders [this is an extension of the scheme in Buckle *et al*. (2018) which only included active proteins]. To identify active protein binding sites we used DNA accessibility measured via DNase-seq experiments, assuming that DNase-hypersensitive sites (DHS) are binding sites (see Methods and Supplemental Methods). Repressive protein binding sites were identified using the chromatin states, with one species binding to H3K9me3 states, and one species representing polycomb repressive complexes and binding to H3K27me3 states. Finally, ChIP-seq for CTCF was used to identify “anchor” sites where loop extrusion is halted. Full details of the simulations are given in Supplemental Methods, and a schematic is shown in Supplemental Fig. S1.

To verify that HiP-HoP gives good predictions for *CCND1*, we generated simulated Hi-C interaction maps and compared them with publicly available data (Rao *et al*., 2014) (Fig. 2A; Supplemental Fig. S1 shows a similar plot but for the entire 3 Mbp region which was simulated, along with the experimental data used as an input). The simulations gave a good prediction of the TAD pattern across the region [clear domains are visible in Fig. 2A; we used the TAD-calling algorithm from the HiCExplorer software (Ramírez *et al*., 2018) to identify boundaries, see Supplemental Fig. S2A-B and Supplemental Methods Section 4]. *CCND1* sits at the far right of a TAD which is bounded by convergent CTCF sites, starts just to the left of *TPCN2*, and ends between *LTO1* and *FGF19*. We shall refer to the three neighbouring domains as the *TPCN2, CCND1*, and *FGF19*-TADs. The gene *MYEOV* sits just left of the centre of the *CCND1*-TAD. More broadly, the dependence of interactions with genomic separation was correctly captured by the model (Supplemental Fig. S2C), and we found good correlation with experiments using several different metrics (*e*.*g*., a Pearson correlation of 0.7 was observed for the directionality index profile, see Supplemental Fig. S2A-F and Supplemental Methods Section 4 for full details).

**Figure 2.**
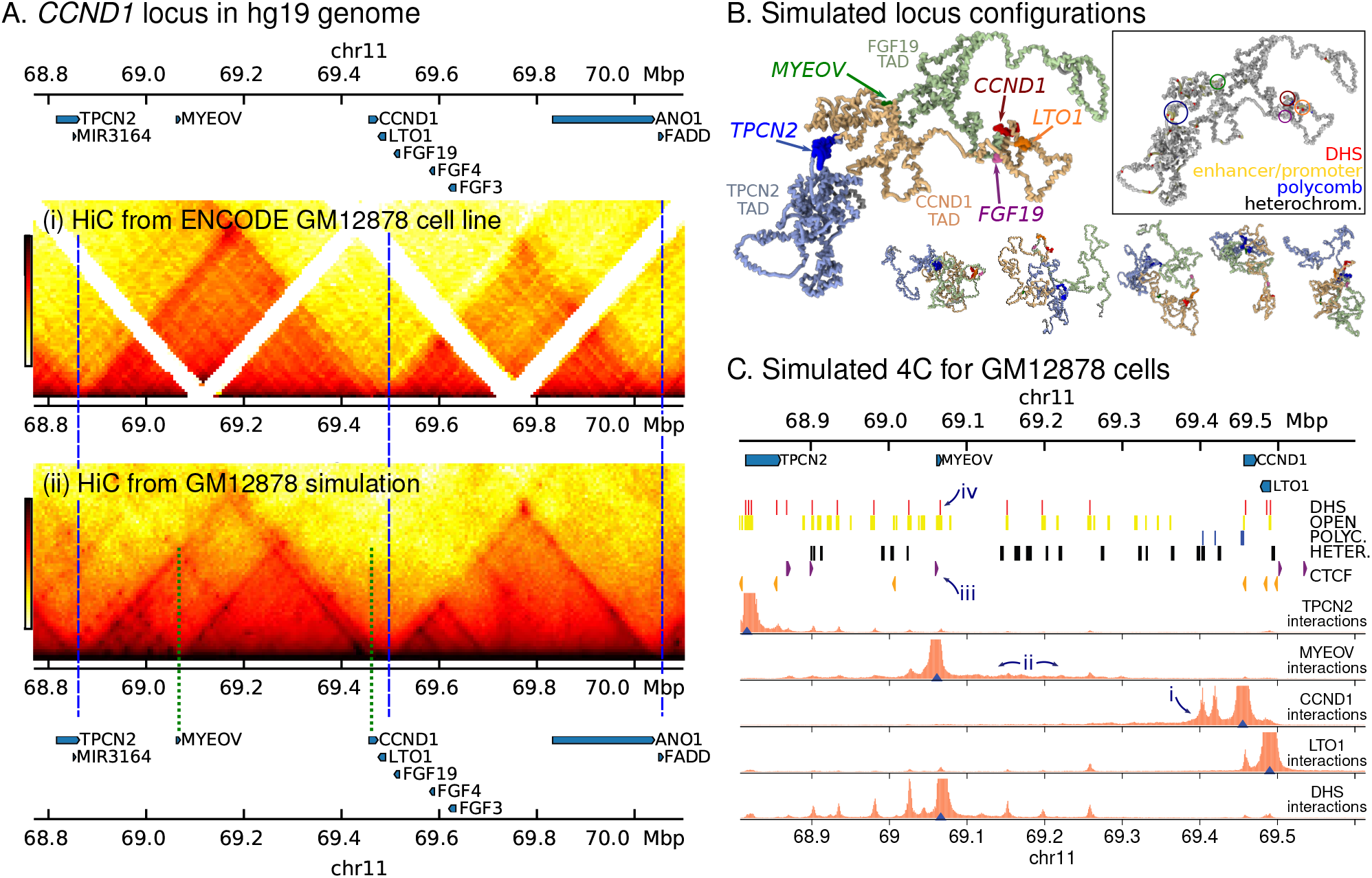
**A**. A map of the genes in the vicinity of *CCND1* is shown above heat maps for: (i) Publicly available Hi-C data for GM12878 cells; and (ii) simulated Hi-C from the same region (obtained from 4400 independent configurations). TAD boundary positions are indicated with blue dashed lines; green dashed lines indicate a “subTAD” region (see text). Some unmappable regions are visible as gaps in the data (white stripes). **B**. Simulation snapshots of the 3D structure of the *CCND1* region, including the *CCND1*-TAD (pale orange) and the two neighbouring TADs (pale green and pale blue; a 1.3 Mbp section of the 3 Mbp region simulated is shown). One representative snapshot is shown large, with five other smaller examples. Several genes are shown in bright colours as indicated. For clarity proteins are not shown. Box: the same configuration is shown but coloured according to the input data used in the simulation, as indicated by label colours; gene positions are circled. **C**. Simulated 4C interaction profiles are shown from four viewpoints [blue triangles; these are positioned at promoters of *TPCN2, MYEOV, CCND1* and *LTO1*, and at a DNase hypersensitive site (DHS) downstream of *MYEOV*]. The height of the orange shaded region represents the relative frequency at which that position interacts with the viewpoint. Data used as simulation input [obtained from ENCODE; see Supplemental Methods and Supplemental Table S3] is shown as coloured blocks above the plots. Red blocks indicate DHS, used to infer binding sites for active proteins. Blue and black blocks indicate regions with chromatin states corresponding to polycomb and heterochromatin respectively, used to infer binding for the corresponding proteins. Yellow blocks indicate regions with chromatin states associated with H3K27ac, and indicate regions which have a more open chromatin structure in the model. Orange and purple arrowheads indicate the position and directionality of CTCF sites, used to infer loop extruder anchors. Some features are labelled with numbered arrows as referred to in the text.

Simulated Hi-C maps tend to show more structure within the domains than experiments (visible as dark spots and stripes; this was also observed in previous HiP-HoP studies). Typically, these features arise in the model due to protein mediated enhancer-promoter interactions. It could be that these interactions are over-estimated in the model, *e*.*g*., because parameters such as the number of bridging proteins are not correct, or because such interactions are disrupted by a mechanism which is not present in the HiP-HoP framework. Another possibility is that the experimental data lacks sufficient resolution to reveal these features. Many of the predicted ‘within-TAD’ interactions are present in publicly available Hi-ChIP and ChIA-PET data which has higher resolution [Supplemental Fig. S2G; promoter-enhancer interactions are also more apparent in targeted 4C (Zhao *et al*., 2006), nucleosome resolution MicroC (Krietenstein *et al*., 2020), and recent Hi-C data which has been treated using a method for removing biases (Lu *et al*., 2020)]. A quantitative analysis of within-domain loops found in Hi-ChIP data shows that the simulations correctly predict an enrichment of these interactions (Supplemental Fig. S2G, and see Supplemental Methods Section 4). In the simulated Hi-C several smaller ‘subTADs’ are also visible; for example, one subTAD is generated by looping interactions between a CTCF site just upstream of *MYEOV* and a CTCF within the *CCND1* gene body [green dashed lines in Fig 2A(ii)]. A typical simulated structure is shown in Fig. 2B; also shown are 5 other representative structures (from a population of 4400 generated).

A more focused view of chromatin interactions at gene promoters is shown in simulated 4C interaction profiles [Fig. 2C; we take a 1 kbp region immediately upstream of a gene’s transcription start site (TSS) to be its promoter]. These reveal that the *CCND1* promoter often interacts with polycomb associated regions (arrow ‘i’ in the figure); this is because there is enrichment of H3K27me3 at the promoter. There are also ‘active’ H3K4me1/3 modifications, which could indicate variation between alleles or across the population, with the gene sometimes expressed and sometimes repressed. Consistent with this, publicly available RNA-seq data (The ENCODE Project Consortium, 2012) shows weak expression of *CCND1* in GM12878 cells (see Supplemental Methods).

To more quantitatively assess the expression of four of the genes within the locus we performed qPCR experiments (see Supplemental Methods and Supplemental Fig. S3). We found that in these cells *TPCN2* and *LTO1* are expressed at slightly lower levels than *CCND1*; in simulations their promoters interact with several nearby DHS. *MYEOV* shows much lower expression than the other three genes (Supplemental Fig. S3A), and is covered by enhancer-associated H3K4me1 and H3K27ac states. While there is no DHS at the *MYEOV* promoter to drive strong interactions, there is a general enrichment of interactions within a broad region downstream of the gene (arrow ‘ii’ in Fig. 2C), driven by the CTCF site immediately upstream of the gene (arrow ‘iii’). Interestingly, there is a DHS just downstream of *MYEOV* (arrow ‘iv’) which interacts strongly with DHS across the locus (including at several promoters, see bottom track in Fig. 2C); this, together with the chromatin states, suggests that this region may have enhancer function.

### A super-enhancer insertion near CCND1 drives changes in 3D structure within the TAD

We next used HiP-HoP to study *CCND1* in the MM cell line U266. These cells possess a genomic rearrangement where an enhancer region from *IGH* is inserted upstream of *CCND1*, within its TAD boundaries. A number of super-enhancers have previously been described in the non-variable *IGH* region, including the long recognised E*α*1, E*α*2 and E*μ* enhancers (Mills *et al*., 1983; Chen and Birshtein, 1997; Mills *et al*., 1997). More recently we proposed that a fourth element E*δ* should be identified as distinct from E*μ*, based on high quality chromatin state mapping (Mikulasova *et al*., 2020b). The ~100 kbp inserted region in U266 includes E*α*1 [accurately mapped by Mikulasova *et al*.(2020b)]. U266 is a useful model because it allows study of the effect of relocating a single *IGH* enhancer within an existing TAD. Strikingly, it leads to significant changes in expression which are similar to those resulting from a reciprocal translocation.

In order to simulate the 3D genome structure in U266 cells we generated an alternative genome build which includes the insert, denoting this hg19_u266 (compared to the unaltered hg19 genome used for GM12878 above). We obtained U266 DNA accessibility and chromatin states from the BLUEPRINT project (Stunnenberg *et al*., 2016), and used CTCF binding site data for B-cells (The ENCODE Project Consortium, 2012). Example configurations are shown in Fig. 3. Figure 4A shows simulated Hi-C results for U266 along with the input data; the insert region is marked in green. For comparison, Fig. 4B shows the intact locus from the GM12878 simulations (no insert).

**Figure 3.**
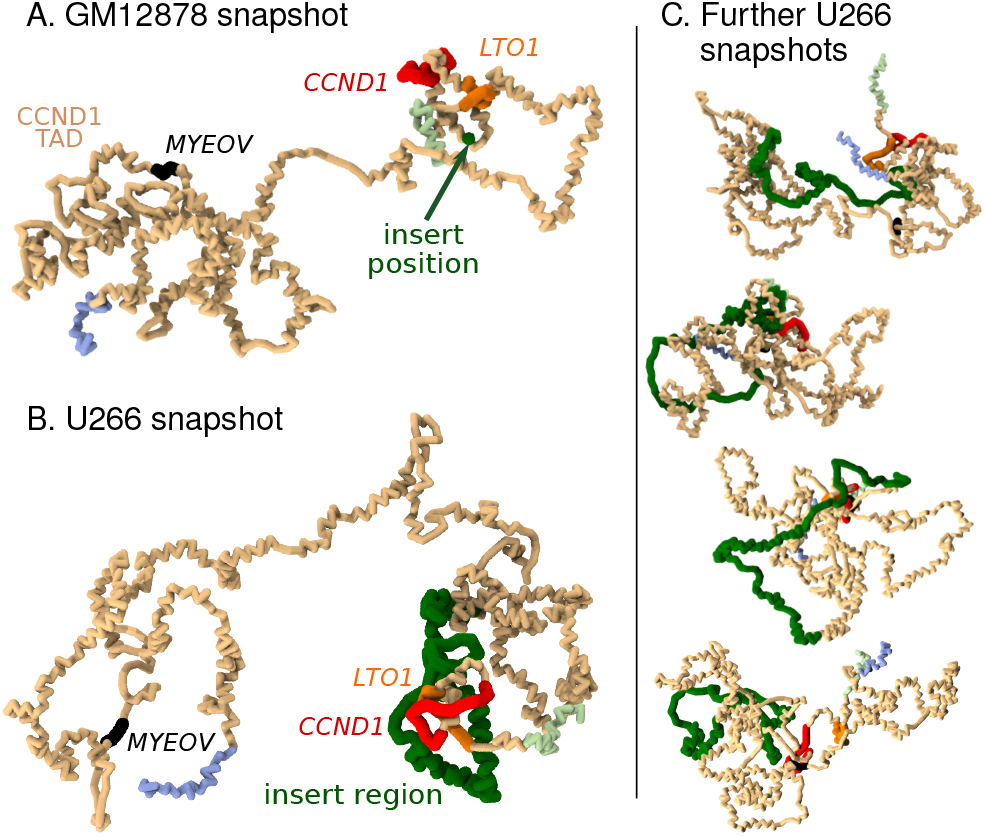
Simulation snapshots. **A**. Example snapshot from simulations showing the *CCND1*-TAD in GM12878 cells (pale orange region). Positions of genes are shown in different colours as labelled. Pale green and pale blue regions are within the neighbouring TADs. **B**. Similar snapshot from U266 simulations. The *IGH* insert containing E*α*1 is shown in green; for comparison, in the snapshot in A the position of the insert is shown as a single green bead. **C**. Four further example snapshots for the U266 cells are shown.

**Figure 4.**
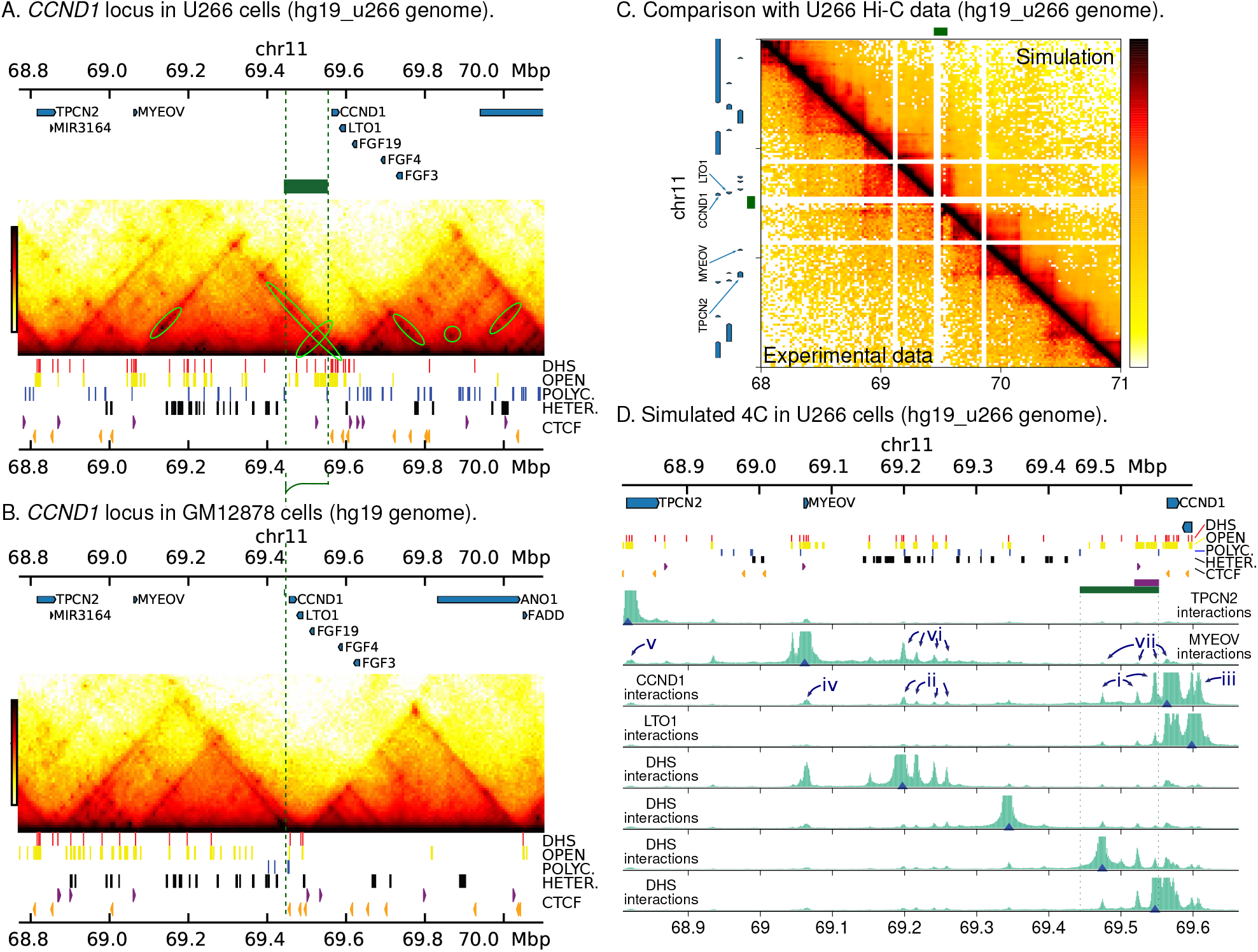
**A**. Simulated Hi-C data is shown below a map of the genes within the rearranged *CCND1* locus as found in the U266 cell line (hg19_u266 genome; a region encompassing the *CCND1* and *FGF19*-TADs is shown). Green rings highlight additional interactions not present in GM12878 cells. The green block and dashed lines show the position of the inserted region. Data used as simulation input [obtained from BLUEPRINT and ENCODE] is shown as coloured blocks under the Hi-C map using the same scheme as in Fig. 2C. **B**. Simulated Hi-C data is shown for the intact *CCND1* locus for GM12878 cells (hg19 genome). The green dashed line indicates the insertion point for the U266 rearrangement. **C**. Side-by-side comparison of simulation and Hi-C data for U266 cells obtained from Wu *et al*. (2017). Experimental data have been aligned to the *in silico* rearranged reference genome hg19_u266 (see Supplemental Methods Section 4 for details). Some unmappable regions are visible as gaps in the data (white stripes); for ease of comparison, the same regions are masked in the simulation map. The lower read depth of the data necessitates a lower map resolution (20 kbp bins compared to 10 kbp bins for GM12878 data). The green blocks to the top and left of the map show the position of the inset region; gene positions are shown to the left. See Supplemental Fig. S4 for a comparison of called TAD boundaries. **D**. Simulated 4C plots are shown for U266 at various viewpoints (blue triangles). In the top four tracks, viewpoints are at promoters of *TPCN2, MYEOV, CCND1* and *LTO1*. The bottom four tracks show reciprocal viewpoints from interacting regions. The green block and dashed lines show the position of the insert, red blocks show DHS, and the purple block shows E*α*1. Some features are labelled with numbered arrows as detailed in the text.

The overall TAD structure is unchanged by the presence of the insert, however, interactions within the domains show significant differences between the two cell lines. Particularly, in U266 there are several strongly interacting regions (dark spots in the Hi-C map, some highlighted with green circles), including close to *CCND1* and within the *IGH* insert itself. Many of these are between DHS, between regions with heterochromatin states, or between regions with polycomb states. DHS often coincide with regulatory elements such as promoters and enhancers; particularly in U266 there is a cluster of DHS near and within *CCND1* which is not present in GM12878. Together these observations suggest that loop extruders and CTCF sites drive TAD formation, while bridging proteins drive interactions within TADs. Extrusion is unchanged by the presence of the insert (which does not contain any CTCF sites), while significant remodelling of chromatin in U266 compared to GM12878 leads to changes within the TAD. To validate this new TAD layout, we compared our simulated interaction map with Hi-C data for U266 cell obtained from Wu *et al*. (2017) (Fig. 4C). Though this data has much lower read coverage than the GM12878 data discussed above, it is sufficient to generate a map at 20 kbp resolution and call TADs (Supplemental Fig. S4); however, there is a region with repetitive DNA sequence within the insert region to which short Hi-C reads cannot be mapped, and this leads to an incorrectly called TAD boundary. Nevertheless, the rest of the TADs in the data match those called from the simulated map, and there is good visual agreement (Supplemental Fig. S4, and see Supplemental Methods Section 4 for details of the analysis). It is clear that *CCND1* is indeed within the domain which encompasses the *IGH* insert, and its promoter shows enriched upstream interactions, validating the simulation predictions. Broader comparison measures also showed significant correlations between the simulation and experimental Hi-C, though the agreement was reduced compared to the GM12878 case (likely due to the lower resolution of the data; Supplemental Fig. S4 and Supplemental Methods Section 4).

Figure 4D shows simulated 4C with viewpoints positioned at gene promoters (top four rows). We found that there were new (compared to GM12878) strong interactions between the *CCND1* promoter and several DHS across the TAD, including three prominent peaks within the insert (arrow ‘i’ in Fig. 4D), several peaks between *MYEOV* and the insert (arrows ‘ii’; some of these DHS were not present in GM12878), and within around the nearby genes (arrows ‘iii’, ‘iv’, and ‘v’). Reciprocal interactions were observed as expected (Fig. 4D, bottom four rows). Many of the regions around the new DHS also gained enhancer or promoter chromatin states compared to GM12878. The new interactions arise because *CCND1* gains DHS and active promoter and enhancer chromatin states at the promoter and across its body (while losing H3K4me1 and H3K27me3 at the promoter). These features are consistent with H3K4me3 broad domains, which have previously been associated with superenhancers (Thibodeau *et al*., 2017; Dhar *et al*., 2018). Their appearance has been implicated in cancer-specific super-enhancer hijacking at a number of oncogenes [including *CCND1*; Mikulasova *et al*. (2020b)].

The promoter of *MYEOV* also gains a DHS in U266 compared to GM12878, and the chromatin state changes such that H3K27ac and H3K4me3 now covers the gene, extending several kbp downstream (another broad domain). It also shows a number of new interaction peaks, particularly in a region downstream where there is a cluster of new DHS with active enhancer and promoter chromatin state (arrows ‘vi’ in Fig. 4D). The *MYEOV* broad domain region as a whole interacts with the *CCND1* broad domain in ~ 3% of configurations, compared to ~ 2% in GM12878. Surprisingly, *MYEOV* shows more frequent interactions with the *CCND1* promoter than with the *IGH* insert, despite the latter being closer genomically (arrows ‘vii’). Taken together these observations suggest that this new *MYEOV* broad domain is active as an enhancer (as noted above the *MYEOV* proximal region also displayed some enhancer-like features in the healthy cell line).

HiP-HoP generates a population of structures (representing a population of cells), and provides full three-dimensional details. We can also therefore measure, for example, separations between specific points or the overall 3D size of a given region. We find that the mean separation of *TPCN2* and *CCND1* tends to be larger in U266 where the insert is present. This is expected because not only is the genomic separation (along the chromosome) of the probes larger, but in the model more of the region has the open (H3K27ac associated) structure. The latter effect is highlighted by examining the 3D size of a region of fixed genomic length around *CCND1* in the two cell lines: the 3D size is larger in U266 cells (Supplemental Fig. S5D, and see also Fig. 6A below).

### The reciprocal translocation t(11;14)(q13;q32) generates an oncogenic TAD fusion

The MCL cell line Z-138 harbours a translocation that dramatically changes the local environment of *CCND1*: it relocates to chromosome 14 where it becomes juxtaposed with the joining region of the *IGH* locus. While the t(11;14) rearrangement is common, the break-point has not previously been mapped in this cell line. Using paired-end read targeted DNA sequencing, we precisely mapped the chromosomal changes (Supplemental Fig. S6). With this information we could then perform simulations of the rearranged genome, which we denote hg19_z138.

Figure 5 shows results from simulations using Z-138 epigenomic data [again obtained from BLUEPRINT, Stunnenberg *et al*. (2016), and ENCODE, The ENCODE Project Consortium (2012)]. The simulated Hi-C map (Fig. 5A) predicts that a new TAD will form, encompassing *CCND1, LTO1* and the *IGH* superenhancers (E*α*2, E*α*1, E*μ*, and E*δ*). The right-hand boundary of the TAD is formed by a pair of divergent CTCF sites, and is conserved from the chromosome 11 boundary observed in GM12878 and other cell types. The left-hand boundary is formed by three inward pointing CTCF sites; a boundary is also seen at this position on chromosome 14 in some other cell types which do not have a translocation [including human umbilical vein endothelial cells, Rao *et al*. (2014)]. In summary, the simulations predict that these two boundaries are conserved after the translocation, and this leads to formation of a new *IGH*-*CCND1* TAD. This new TAD has a length of about 500 kbp, about 20% smaller than the ∼ 640 kbp *CCND1*-TAD observed in GM12878 cells (the 3D volume of the TAD in Z-138 is also correspondingly smaller). Interestingly, one can also see a region within the TAD which shows stronger enrichment of iterations (a ‘subTAD’ of length 407 kbp, Fig. 5B); this arises due to looping between a CTCF site just upstream of E*α*2 and a CTCF within *CCND1*. The latter is the same CTCF site which forms a subTAD in GM12878.

**Figure 5.**
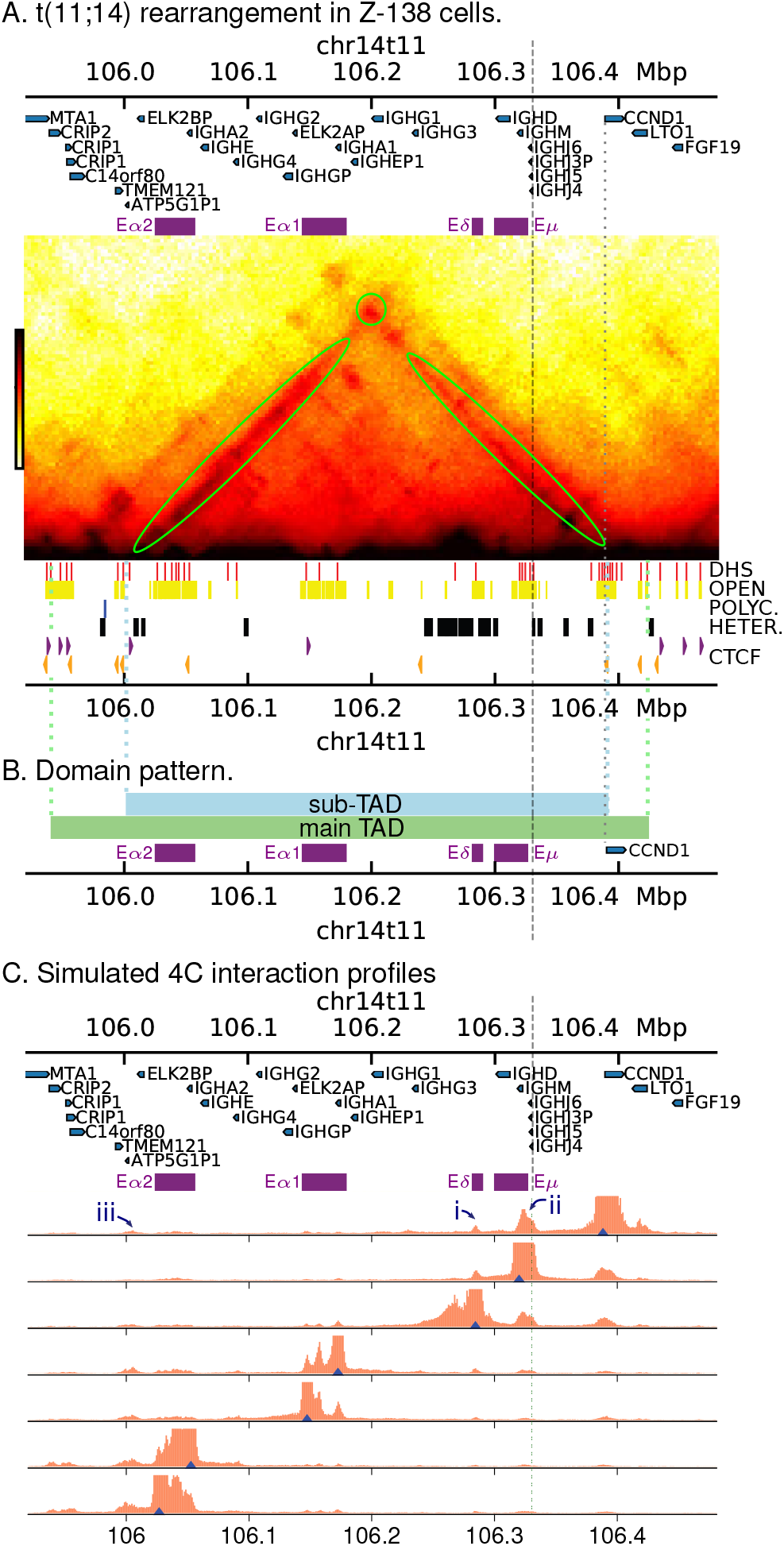
**A**. Simulated Hi-C for Z-138 cells which possess a t(11;14) translocation (hg19_z138 genome build). Positions of the *IGH* superenhancers are shown as purple blocks above the map. Simulation input data (DHS, chromatin states and CTCF peaks) are shown under the map as in previous figures. The grey dashed line shows the position of the translocation, while the grey dotted line shows the position of the *CCND1* promoter. **B**. Coloured blocks show the domain pattern. The green block shown the domain obtained via a TAD-calling algorithm (see Supplemental Methods), while the blue block shows a subTAD region which is visible in the interaction map in A. The subTAD boundary positions are determined by the positions of CTCF sites. Coloured dotted lines show where the boundaries lie on the map in A. **C**. Simulated 4C from viewpoints positioned at DHS across the region (blue triangles). Some features are highlighted with numbered arrows as discussed in the text.

Figure 5C shows simulated 4C with viewpoints at *CCND1* and several positions within the translocated *IGH* super-enhancers. *CCND1* shows prominent interaction peaks in both E*δ* and E*μ* (arrows ‘i’ and ‘ii’ in the figure; the interaction is strongest where several DHS are clustered together). There are also smaller interaction peaks within E*α*1 and E*α*2, and at a number of other DHS (notably a cluster to the left of E*α*2, arrow ‘iii’). Similar to U266, there are a number of DHS at the *CCND1* promoter and within the gene body which are not present in the GM12878 cell line. In the simulations the interactions of the promoter are driven by binding (and molecular bridging) of active proteins; this bridging is likely further promoted by CTCF/cohesin driven loop extrusion. Viewpoints positioned at DHS within the enhancers show reciprocal interactions with *CCND1*.

Here we have simulated the break-point and rearrangement as mapped for the Z-138 cell line (about 60 kbp upstream of *CCND1*). It is also possible to simulate alternative rearrangements to better understand the effect of break-point variation between patients. Up to 50% of t(11;14) MCL break-points are within the so-called ‘major translocation cluster’ about 120 kbp upstream of *CCND1* (Jares and Campo, 2008), with the rest scattered throughout a broader 380 kbp region around this. Supplemental Figure S7 shows simulated 4C from two different examples where the chromosome 14 break-point is kept the same but we move the break-point on chromosome 11 to different positions between 11 and 100 kbp upstream of *CCND1*. We found that in general, interactions between the gene and the E*μ* enhancer are weaker if their genomic separation is larger.

### Chromatin remodelling and *CCND1* 3D structures differ in U266 and Z-138

The changes to chromatin states and DNA accessibility at *CCND1* differ subtly between the two cancer cell lines. This is visible in Figs. 4A and 5A, but is more evident if the data are mapped back to the hg19 reference genome (Supplemental Fig. S8 shows the chromatin states and DHS at three regions within the *CCND1* locus where there is significant remodelling compared to GM12878). In both U266 and Z-138 the *CCND1* gene body gains several DHS and a H3K4me3 broad domain appears. From 10 distinct DHS within *CCND1*, 5 are common to both cancer cell lines, 3 are specific to Z-138 and 2 are specific to U266. In other words, the different rearrangements lead to similar, but not identical, changes to the chromatin structure at *CCND1*. In U266 cells, two other regions within the TAD also show substantial changes: a region covering and extending downstream of *MYEOV* and a region between *MYEOV* and *CCND1* (location of arrows ‘ii’ and ‘vi’ in Fig. 4D) both gain a H3K4me3 broad domain and several new DHS. We note that the pattern of chromatin states and accessibility at the *IGH* enhancers also varies between the U266 and Z-138 cell lines (see Supplemental Fig. S9).

The changes to the chromatin states at *CCND1* has a striking effect on the 3D structure of the gene: in Fig. 6A we plot the distribution of the simulated 3D size of the gene body (quantified by the radius of gyration, *R*_*g*_, see Supplemental Methods). In GM12878 the gene is on average more compact than in the other cell lines. The snapshots in Fig. 6A show typical configurations: the differences in the size of the gene, the chromatin states, and the DHS pattern are clear. In GM12878 the gene has a crumpled structure, while in U266 it is more stretched out.

**Figure 6.**
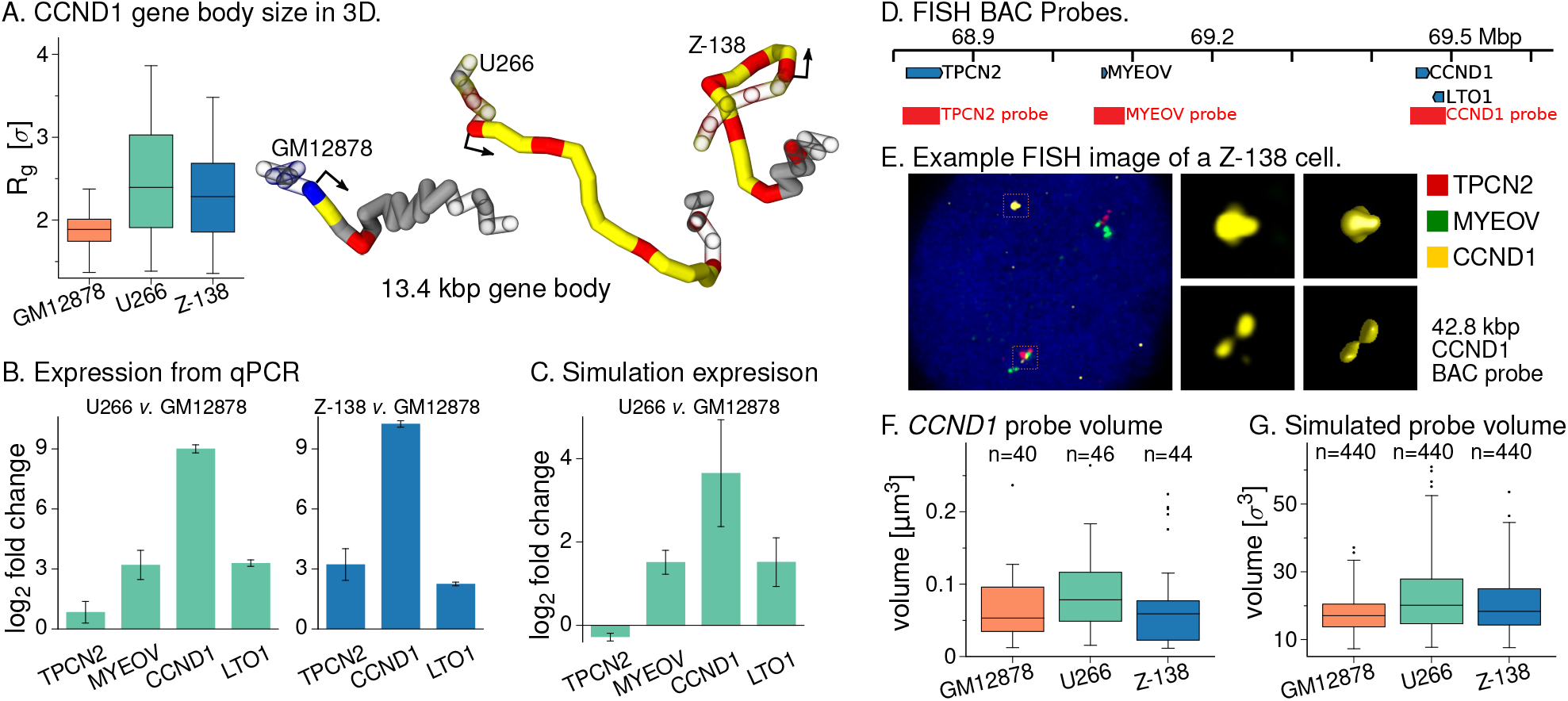
**A**. Left: box plot showing the distribution of the radius of gyration of the *CCND1* gene body (a measure of its 3D size) in simulations for each of the three cell types (440 configurations for each case). Right: Typical snapshots of the gene and a 5 kbp flaking region on each side. The gene body is shown as a solid tube, with flanking regions shown as an outline. The arrow shows the transcription start site. The polymer is coloured according to the simulation input data: red indicates active protein binding sites (DHS), blue polycomb protein binding sites, and yellow are open chromatin (H3K27ac). **B**. Left: bar plot showing the change in expression level in U266 cells compared to GM12878 obtained from qPCR measurements (see Supplemental Methods Section 6 and Supplemental Fig. S3 for further details). The bar height gives the log_2_ fold change based on an average over three replicates; the error bar shows the standard error in the mean. Right: similar plot for Z-138 cells compared to GM12878. No *MYEOV* expression was detected in these cells. **C**. Bar plot showing the fold change in expression level in U266 compared to GM12878 based on the expression prediction from simulations. As detailed in the text, we do not expect a linear mapping between the predicted and real expression levels, so one can only compare qualitatively with the qPCR. Nevertheless, this shows the same trend across the genes as in panel B. See Supplemental Methods for further details. **D**. Map of the *CCND1* locus (hg19) showing the positions of the fosmid probes used in FISH experiments. **E**. Example FISH images. The left image shows a whole cell with red, green and yellow probes covering different genes as indicated. Middle images show the yellow probe spots only (regions indicated by orange boxes in the whole-cell image). Right images show a 3D reconstruction of the gold probes as used to measure the volume (see Supplemental Methods for details). **F**. Box plot showing the distributions of the volume of the probe covering *CCND1* in the three cell lines (see text and Supplemental Methods for details). The number of probe spots measured in each case (*n*) is indicated. See Supplemental Fig. S10 for similar plots for the other probes. **G**. Box plot showing a similar measurement but extracted from *n* = 440 simulated configurations from each cell line (see Supplemental Methods for details of the calculation; simulation length units roughly map to physical units as *σ* ≈ 21.8 nm, so *σ*^3^ ≈ 1.04 × 10^−5^µm).

Protein mediated loops can form between the DHS within the gene body which reduce its 3D size: variation in the number of such loops present at any one time leads to the large variation in *R*_*g*_. The snapshot for the Z-138 case shows a configuration where a loop forms between a DHS within the gene body and a DHS in the upstream region. Since there are more DHS across the gene in Z-138 there are more possibilities for loops to form: smaller *R*_*g*_ values are therefore more likely and there is less variation.

To see how this large change in 3D structure correlates with gene expression, we performed additional qPCR experiments comparing the expression of the four genes in the locus between the cell lines. Figure 6B shows their fold change in expression in U266 compared to GM12878 (see also Supplemental Fig. S3B). Notably, while all of the genes show an increase, that of *CCND1* is by far the largest: more than 500-fold. The next largest change is for *LTO1* which shows a ~ 9.8-fold increase. We considered whether these changes could be predicted from the simulated structures. In our simulations the active proteins represent complexes of transcription factors and polymerases, and one might therefore associate binding of these to a gene promoter site with transcription. Indeed, in previous work using a simpler model we showed that the fraction of time a promoter is bound by an active protein during the simulation is correlated with gene expression measured experimentally via GRO-seq (Brackley *et al*., 2020). In the simulations the active proteins can bind at promoters which overlap a DHS; they can also simultaneously bind multiple DHS to form molecular bridges. An emerging feature of the model is that clusters of proteins form at positions where there are bridges between DHS, and that in most cases where a promoter is bound by a protein, it belongs to one of these clusters. This means that the likelihood that a promoter is bound by a protein depends on the local 3D structure, consistent with a mechanism where enhancer-promoter interactions and/or formation of transcriptional protein condensates drives expression. Here we calculated the fraction of simulated configurations where a given promoter is bound by a protein. This measure does not take into account enhancer ‘strength’ or compatibility, and can only have a value between 0 and 1; therefore, we only expect a correlation and not a quantitative mapping to expression. Nevertheless, the predictions are largely consistent with the qPCR data, even if this is not a direct validation of the predicted structures (see Supplemental Methods and Supplemental Fig. S3Ei for further details). An even better correlation between simulations and data is obtained if, rather than considering protein binding, we instead ask how often the promoter interacts with (forms loops to) chromatin which has an enhancer associated state (Fig. 6C). We note that there are simpler models which aim to predict expression, for example based on histone modification levels near gene promoters (Fulco *et al*., 2019), but these do not provide the same insight and context revealed by the predicted 3D structures.

In Z-138 cells, *CCND1* expression shows an even larger 1,200fold increase compared to GM12878 (qPCR data shown in Supplemental Fig. S3C); *LTO1* and *TPCN2* also show increased expression in Z-138 compared to GM12878 cells, but we did not detect *MYEOV* expression. Only *CCND1* and *LTO1* were present within the region simulated, and simulations correctly predicted the former would display the greater fold change compared to GM12878 cells (Supplemental Fig. S3Eii).

To verify that the change in 3D gene size is observed *in vivo*, we performed fluorescence *in situ* hybridisation (FISH) microscopy using fosmid probes [see Methods and Supplemental Methods]. We note that the probes cover a region several times larger than the *CCND1* gene body (42.8 kbp compared to the 13.4 kbp gene), so this is an indirect measure. Figure 6E shows a typical image from the Z-138 cells. First, we note that the positioning of the spots is consistent with the translocation. One of each of the red, green and gold spots are together in a group (on the non-translocated chromosome 11); the other gold (*CCND1*) probe is alone, while the other red and green probes (*TPCN2* and *MYEOV* respectively) are together. From the images we obtain the 3D volumes of the probes; a similar measure is obtained from the simulations. The simulations predict that compared to the GM12878 cell line, the volume occupied by the *CCND1* probe region will be on average 30% larger in U266 cells and on average 16% larger in Z-138 cells. They also predict that the *MYEOV* probe volume will be on average 36% larger in U266 cells compared to GM12878, while the *TPCN2* probe will be the same volume in both of these cell lines. Experimentally, we obtained measurements from at least 20 cells for each cell line (with two spot measurements in each; the numbers of cells where spots were clearly visible was limited by some technical aspects, see Supplemental Methods). The data showed trends which were generally consistent with the simulations: the *CCND1* probe volume was on average 30% larger in U266 compared to GM12878, while the *MYEOV* probe was on average 23% larger. However, in the experiments the differences were not statistically significant; nevertheless, the simulations are still consistent with this, as taking a random subset of the simulation measurements such that there were the same number as in the experiments also did not show any significant differences between the probe volumes in the different cell lines (see Supplemental Methods Section 7 for further details). Plots showing the distributions of the *CCND1* probe volumes for experiments and simulations are shown in Figs. 6F and G (Supplemental Fig. S10 shows similar plots for the other probes). We note that while the simulations are consistent with the *in vivo* trends, we would not expect the model to be able to predict quantitative differences in probe volumes; this is consistent with previous work where predictions for 3C-based data (interactions) were better than those for microscopy data (Buckle *et al*., 2018).

### The oncogene structure of *CCND1* is driven by chromatin remodelling

We have observed that in these cell lines, the genome rearrangement is accompanied by extensive chromatin remodelling. In particular, in both U266 and Z-138 a H3K4me3 broad domain containing several DHS appears over *CCND1*. A likely scenario is that after the rearrangement, the *IGH* super-enhancers recruit transcription factors, chromatin remodellers, and architectural proteins, to the region. The resulting changes then in turn disrupt the wider 3D structure, leading to dysregulation of *CCND1*. The HiP-HoP computational framework provides a unique opportunity to examine these two effects in isolation: we can rearrange the genome without otherwise changing the chromatin states, or we can use chromatin states from a cancer cell line without including any genomic rearrangement. Importantly, it is not possible to do this *in vivo*, and these *in silico* situations cannot be engineered in reality. Nevertheless, they provide insight.

Figure 7A shows results from a simulation where chromatin state and DHS data for GM12878 cells were used as input, but the hg19_u266 genomic insert rearrangement was included (as depicted in the cartoon). In Fig. 7C we compare 4C results from GM12878 simulations with and without the insert; this shows that adding the insert has only a small effect on the interaction profile of the gene promoters. The profiles for *TPCN2* and *MYEOV* are unchanged. Also, despite the insert containing a DHS within a H3K27ac region, there is little interaction with the nearby *CCND1* promoter (which has a polycomb chromatin state). On the other hand, there is some interaction between the insert and *LTO1*. Using the simulations to predict gene expression as above, these results suggest that without any subsequent chromatin remodelling, the presence of the insert alone would not lead to *CCND1* up-regulation. This is in contrast to the measured change in RNA levels via qPCR (Supplemental Fig. S3B-C), which show approximately 500-fold and 1200-fold increases in *CCND1* expression in U266 and Z-138 respectively, compared to GM12878.

**Figure 7.**
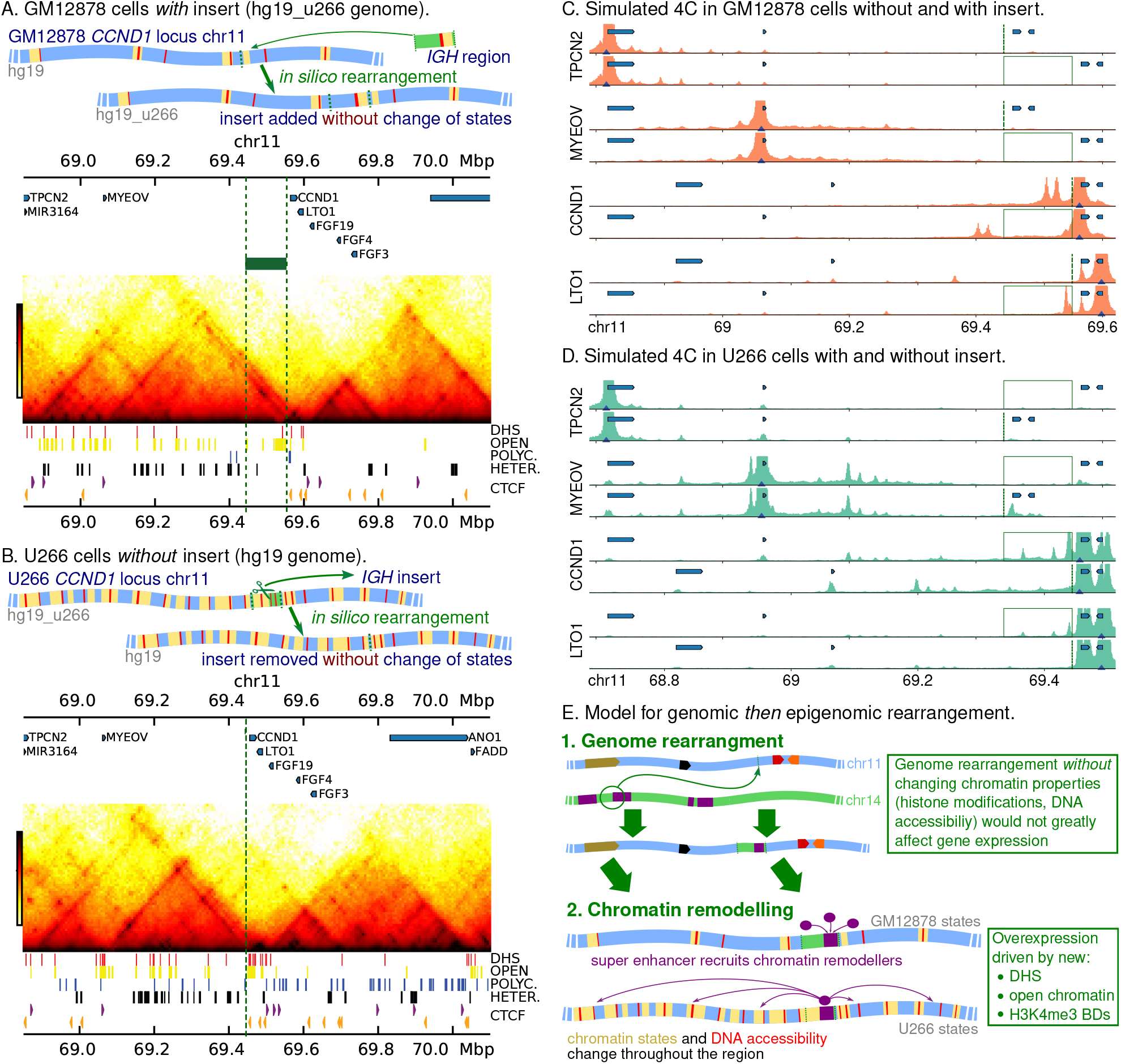
**A**. Top: cartoon showing an *in silico* genome rearrangement where an *IGH* region is inserted into the GM12878 *CCND1* locus, and the GM12878 (healthy) chromatin states are retained (*i*.*e*., the hg19_u266 reference genome is used). Bottom: Hi-C map generated from simulations with this *in silico* rearrangement. The input data are shown below the map; the green block and dashed lines show the position of the insert. **B**. Top: cartoon showing an *in silico* genome rearrangement where the *IGH* insert is removed from the U266 *CCND1* locus, and the U266 chromatin states are retained (*i*.*e*., the hg19 reference genome is used). Bottom: simulated Hi-C generated from simulations with this *in silico* rearrangement. The green dashed line shows where the insert has been removed. **C**. Simulated 4C are shown for simulations using GM12878 chromatin states both with and without the insert (position indicated with green lines). Positions of genes are shown by blue blocks (from left to right, *TPCN2, MYEOV, CCND1* and *LTO1*); **D**. Simulated 4C are shown for simulations using U266 chromatin states both with and without the insert. **E**. Cartoon showing a model where first the genomic rearrangement brings the *IGH* super-enhancer into the *CCND1* TAD; this then leads to recruitment of chromatin remodellers *etc*. to the region and local chromatin states and DNA accessibility are altered. This could be described as an “epigenomic rearrangement”.

Figures 7B and D show similar results from simulations where U266 epigenomic input data is used with the intact hg19 genome (no insert, as shown in the cartoon). Here we see the effect of the chromatin remodelling in the absence of the insert. We find that in this case the *CCND1* promoter still shows interaction peaks across the *CCND1* gene body, at regions downstream (in and around *LTO1*), and to a lesser extent the H3K4me3 broad domain regions upstream. Beyond the loss of interactions with the insert itself, the main effect of removing the insert is an increase in interactions between *CCND1* and upstream DHS, consistent with their reduced genomic separation. The simulations also predict that very little change in gene expression level would result from removing the insert while keeping the U266 chromatin states. Together these results suggest that it is chromatin remodelling which drives the changes in the 3D structure of the locus in terms of promoter-enhancer interactions, and this in turn drives up-regulation of cyclin D1.

As well as simulating different rearranged genomes, it is also possible to directly edit the input data, *i*.*e*., we can edit the epigenome *in silico*. For example, we found little change in *CCND1* interactions if we take chromatin states from GM12878 cells for chr11, and add an insert from chr14 with states from the U266 cells (Supplemental Fig. S11A-B). In a simulation of the U266 cells with the insert where we “switched off” protein binding at four DHS within the promoter and gene body of *CCND1*, there was a loss of *CCND1* interactions compared to the previous U266 simulations (Supplemental Figs. S11C-E).

## Discussion

In this work we have adapted the HiP-HoP simulation model to study chromatin 3D structure and the effect of genome rearrangements in malignant and non-malignant B-cells. HiP-HoP simulations predict 3D structures from DNA accessibility, chromatin states, and CTCF binding data. We used these structures to simulate Hi-C and 4C data, and to make single cell-like measurements. Importantly, Hi-C data are *not* used as an input, so in this sense the model is truly predictive for 3D structures. Here, by “rearranging” the input data we generated predictions for the effect of genome rearrangements which are found in MCL and MM cell lines.

We first confirmed that the HiP-HoP model gives good predictions for the *CCND1* gene locus by performing simulations of the healthy B-cell derived GM12878 lymphoblastoid cell line. The TAD pattern observed in Hi-C (Rao *et al*., 2014) is clearly reproduced by the simulations, with boundaries at CTCF sites. In these cells *CCND1* is only expressed at very low levels; the promoter region has a polycomb associated chromatin state, and our simulations predicted interactions between the promoter and other polycomb regions. There are several parameters in our model, and varying these could affect the quality of the predictions; however, our previous work (Buckle *et al*., 2018) showed that variation of most parameters has a small effect on the predicted interaction, and this is mostly quantitative rather than qualitative (*e*.*g*., increasing the number of simulated proteins tends to increase chromatin interactions, but without changing the positions of peaks). Our parameter choice here was based on those in Buckle *et al*. (2018) (though due to the different set up, we could not preserve all parameters, see Supplemental Methods), and it may be that predictions could be improved by optimising the parameters for this specific locus.

Further simulations predicted that the TAD structure around *CCND1* is preserved in the U266 MM cell line which possesses a rearrangement where a super-enhancer from the *IGH* locus is inserted upstream of *CCND1*. This is consistent with previously published low-resolution Hi-C data in that cell line (Wu *et al*., 2017). In these cells the chromatin around *CCND1* is remodelled: several new DHS are established, and a broad region gains an active promoter chromatin state (H3K4me3 and H3K27ac). The simulations predicted that the *CCND1* promoter interacts with these new DHS in the gene body, as well as with a number of DHS within the inserted *IGH* region and the neighbouring gene *LTO1* (interactions which are not present in GM12878 cells).

We also performed simulations of the MCL cell line Z-138, where *CCND1* translocates with chromosome 14 becoming juxtaposed with the *IGH* E*μ* enhancer. In this cell line a TAD boundary formed by a cluster of CTCF binding sites downstream of *CCND1* was preserved after the translocation, as was a boundary to the left of the *IGH* super-enhancers (presumably in healthy cells this isolates the super-enhancers from other nearby genes). In other words, the simulations predict a fusion of the *CCND1* and *IGH* regions into a new TAD – an oncogenic TAD fusion (Valton and Dekker, 2016). In this new arrangement the promoters of *CCND1* and *LTO1* readily interact with each other and with the proximate E*μ* and E*δ* super-enhancers, while weaker interactions were observed with the more distant E*α*1 and E*α*2.

*In vivo*, genomic rearrangements are accompanied by chromatin remodelling; within our simulation scheme it was also possible to examine the effect of a rearrangement in the absence of remodelling. Simulations of GM12878 cells with the E*α*1 enhancer DNA inserted *in silico* upstream of *CCND1* (with GM12878 chromatin states otherwise unchanged) showed only very minor changes in terms of 3D structure compared with the unaltered genome. We also performed a simulation using U266 chromatin states, but without the insert (*i*.*e*., the epigenomic rearrangement is included, but not the genomic rearrangement, an experiment that is only possible to do *in silico*). This showed little change compared to the U266 case with the insert. Together this suggests that it is the local chromatin remodelling, rather than the proximity of *CCND1* to the *IGH* enhancers *per se*, which drives gene deregulation. Or in other words, a genomic translocation leads to an epigenomic translocation, which drives cyclin D1 overexpression. The remodelling includes the appearance of an H3K4me3 broad domain over *CCND1*; the results here support our previous work suggesting that such broad domains are associated with super-enhancer hijacking (Mikulasova *et al*., 2020b). In U266 two other nearby regions become H3K4me3 broad domains, one covering the gene *MYEOV* (this region also appears to have some enhancer-like properties in the healthy cell line). Importantly, the chromosomal locations which are predicted to strongly interact with *CCND1* could be used for targeting in future experiments which aim to uncover the mechanisms through which broad domains are generated. For example, one could ask whether using CRISPR-dCas9 to tether KRAB or another enzyme at the U266 insert is sufficient to reverse the nearby epigenetic changes.

In principle, our method can be applied to simulate any genome rearrangement for which input data is available. One current limitation is that the input data comes from both alleles of the locus, so for example in the Z-138 case the chromatin states are based on a combination of histone modifications from the translocated and non-translocated *CCND1* loci. We note that this is a limitation of the input data rather than the simulation scheme; if the full sequence of a given cell line or patient sample were available, ChIP-seq reads could be aligned in an allele specific way. It may also be possible to use the simulations to explore hypotheses about differences between alleles, *e*.*g*., if one were to assume one copy of the locus has active chromatin states and the other inactive states, how does the 3D structure differ? A simplifying assumption in the model is that all DHS are binding sites for a general active factor, and it would be interesting to incorporate more detail in the future, *e*.*g*., if transcription factors specific to a particular oncogene are known (Brackley *et al*., 2016a). Other possible improvements to the model include adding further “compaction levels” in the heteromorphic polymer (which might improve microscopy predictions), or adding protein-protein interactions promoting larger liquid-like phase separated protein droplet formation [augmenting the small protein clusters which form via bridging-induced attraction in the present model; Brackley *et al*. (2013)]. There are also some more general limitations to our approach. We have focussed here on translocations which reposition superenhancers; it is unclear what insight could be gained in other situations (e.g., gene fusions, or frameshift mutations). We have not yet considered copy number variations and gene dosage, and have assumed that we are simulating an initial driver rearrangement; using the approach to understand the evolution of genomic abnormalities which arise sequentially would be more challenging.

It would be of interest in the future to also simulate the *dynamics* of histone modification and DNA accessibility; as noted above, our simulations imply that the genomic rearrangement and repositioning of super-enhancers drives local changes to these chromatin features. For example, including histone modifying enzymes explicitly in a model such as HiP-HoP would open the possibility to understand this process, and to predict how a rearrangement changes the 1D chromatin properties as well as the 3D structure. In the long term, it may then be possible to predict changes to expression using only input data from healthy cells, plus the coordinates of the rearrangement.

In summary, our work strongly suggests that genome rearrangements drive a subsequent epigenomic rearrangement, which in turn leads to deregulation and proto-oncogene activation. We have demonstrated that polymer physics-based modelling can be useful for understanding the structural consequences of genome rearrangements, and can help to focus future experimental work. It would be interesting to see if in the future, such simulations could also help us understand the mechanisms behind the epigenomic rearrangement. This would clearly be important for the development of any therapies which seek to interfere with that process.

## Methods

### HiP-HoP simulations

We used the HiP-HoP model as detailed in Buckle *et al*. (2018); full details are given in Supplemental Methods. In brief, chromatin is represented as a chain of beads (each representing a 1 kbp region), and we evolve the configuration of this chain using a molecular dynamics scheme and the LAMMPS simulation software (Plimpton, 1995). To improve efficiency, in each simulation we include 40 Mbp of chromatin (40,000 beads) which includes 11 copies of the region of interest; for each region we perform two such simulations to generate 4400 individual configurations each of which can be said to represent a single cell. The chromatin concentration is roughly matched to that of a typical nucleus. From the configurations we generate simulated Hi-C and 4C and single cell-like measurements. As detailed in the text, three different mechanisms are included to drive the chain into specific structures: diffusing bridge forming proteins, loop extrusion, and a heteromorphic polymer structure. DNase hypersensitive sites (DHS) are used to identify binding sites for an active protein (we use the simplifying assumption that all DHS are the same and bind this protein, which represents a general complex of polymerase and transcription factors). We use chromatin state data to identify binding sites for two species of repressive protein (*e*.*g*., representing HP1 and polycomb repressive complexes), and to identify regions which have an open chromatin structure. ChIP-seq data for CTCF is used to identify direction-dependent anchor sites for loop extrusion. Full details of the input data treatment are given in the Supplemental Methods. Details of publicly available data used for simulation input and validation are given in Supplemental Table S3 and Supplemental Methods.

### Quantification of gene expression

For gene expression measurements, total RNA was extracted using RNeasy kit (QIAGEN, D usseldorf, Germany) from between 5 ×10^6^ and 10 × 10^6^ Z-138, U266 and GM12878 cells. For qPCR, cDNA was synthesised from 1µg of total RNA using random primers (Promega, Wisconsin, USA) then treated with M-MLV Reverse Transcriptase (Promega, Wisconsin, USA). 2µg cDNA reaction were amplified with SYBR green master mix (Invitrogen, Massachusetts, USA), 10µM forward and reverse primers for *LTO1, TPCN2, CCND1* and *MYEOV* (Sigma-Aldrich, Missouri, USA) with 40 cycles of PCR (95°C for 15 s, 60°C for 60 s) after initial denaturation (95°C for 10 min). Cycle threshold values were normalized by comparison with *GAPDH* expression. See Supplemental Methods for further details (oligonucleotide sequences are shown in Supplemental Table S2).

### Genome rearrangement breakpoint mapping

Breakpoint mapping in U266 and Z-138 cells was performed as described previously (Mikulasova *et al*., 2020b,a); further details are given in Supplemental Methods.

### FISH analysis

For FISH microscopy, briefly, fosmid clones found to cover *TPCN2* (G248P87917D11/WI2-1721H21), *MYEOV* (G248P87014D2/WI2-2222G4) and *CCND1* (G248P86668E2/WI2-2191I3) loci were obtained from BACPAC resources (California, USA). DNA was fluorescently labelled by nick translation. Probes and cell line DNA were denatured at 75°C for five minutes followed by hybridisation at 37°C overnight. Further details are given in Supplemental Methods.

## Data access

All raw and processed sequencing data generated in this study have been submitted to the NCBI BioProject database (https://www.ncbi.nlm.nih.gov/bioproject) under accession number PRJNA635269 (release pending). The qPCR, FISH and simulation data generated in this study have been submitted to the Edinburgh DataShare repository (https://datashare.ed.ac.uk) accession number pending.

## Acknowledgements

Work in the DR lab is supported by a Wellcome Trust Seed Award in Science (206103/Z/17/Z). BMJ acknowledges funding from FEDER / Ministry of Science and Innovation - Spanish State Research Agency under the project RTI2018-094788-A-I00, and La Caixa Banking Foundation under the r project LCF/BQ/PI19/11690001. Work in the LJR lab was supported by CCLG Little Princess Trust (NK) and an MRC Di- MeN DTP studentship (DK). CAB acknowledges support from European Research Council (ERC CoG 648050 THREEDCELLPHYSICS).

## Author contributions

AM and BAW performed the Z-138 break-point mapping and sequencing; DK performed bioinformatics analysis and cell line qPCR experiments. NK performed cell line qPCR experiments, FISH probe generation and post capture analysis of images. LJR performed FISH probe generation, hybridisation and post capture analysis of images. RB-P performed confocal microscopy and post-capture deconvolution and image analysis. CAB designed and performed the simulations. DR, LJR and CAB conceived and designed the research, interpreted the results, and wrote the paper. All authors contributed to writing the paper.

