## Supplemental Material for "High-resolution simulations of chromatin folding at genomic rearrangements in malignant B-cells provide mechanistic insights on proto-oncogene deregulation"

[1] *Biosciences Institute, Newcastle University, Newcastle upon Tyne NE2 4HH, UK*
[2] *Translational and Clinical Research Institute, Newcastle University, Newcastle upon Tyne NE2 4HH, UK*
[3] *Bio-Imaging Unit, Newcastle University, Newcastle upon Tyne NE2 4HH, UK*
[4] *Melvin and Bren Simon Comprehensive Cancer Center, Division of Hematology Oncology, Indiana University, Indianapolis, IN, USA*
[5] *Josep Carreras Leukaemia Research Institute (IJC), IJC Building, Campus ICO-Germans Trias i Pujol, Ctra de Can Ruti, Camí de les Escoles s/n, 08916 Badalona, Barcelona, Spain*
[6] *SUPA, School of Physics and Astronomy, University of Edinburgh, Peter Guthrie Tait Road, Edinburgh EH9 3FD, UK*

#### Supplemental Material

**2 Simulation input data** 3

**3 Simulation Measurements** 4

**4 Chromosome-conformation-capture data for comparison with simulations** 4

**5 Cell lines** 6

**6 Gene expression measurements** 6

**7 Fluorescence *in situ* hybridisation microscopy** 6

**8 Break-point mapping in U266 and Z-138 cells** 7

**9 Supplemental Tables** 9
Table 1. Chromatin states . . . 9
Table 2. Oligonucleotide sequences used in qPCR experiments . . . 9
Table 3. Data sources . . . 9
Table 4. Data availability . . . 9

**10 Supplemental Figures** 10

#### 1 Simulation Methods

We use the HiP-HoP model to perform simulations of a chromatin region as detailed in Buckle *et al.* (2018). In this scheme, a section of a chromatin fibre is represented as a chain of beads connected by springs; this is a common model from polymer physics (Barbieri *et al.*, 2012; Brackley *et al.*, 2013; Fudenberg *et al.*, 2016). We then use molecular dynamics simulations to evolve the configuration of this polymer based on a set of phenomenological potentials which describe how the beads interact. Each bead represents a 1 kbp chromatin region. Three additional model ingredients drive the chromatin configuration: interactions with spheres representing complexes of proteins (Brackley *et al.*, 2013, 2016, 2017; Brackley, 2020), a heteromorphic polymer structure, and loop extrusion (Fudenberg *et al.*, 2016; Sanborn *et al.*, 2015). The original HiP-HoP model includes a single species of model proteins which bind to "active sites" on the polymer. In the present work we extend this scheme by adding two additional protein species which bind different repressed chromatin regions.

#### 1.1 The polymer model

Interactions between the polymer beads are defined by four potentials. First, non-adjacent beads interact sterically via the Weeks-Chandler-Anderson (WCA) potential

$$V_{\mathrm{WCA}}(r)=\begin{cases} 4k_BT\left[\left(\frac{\sigma}{r}\right)^{12}-\left(\frac{\sigma}{r}\right)^{6}\right]+k_BT & r\le 2^{1/6}\sigma,\\ 0 & \text{otherwise,}\end{cases} \qquad (1)$$

where $r$ is the separation of the beads, $T$ and $k_B$ are the temperature of the system and the Boltzmann constant, and $\sigma$ is the bead diameter. Second, adjacent beads are connected by finitely-extensible non-linear elastic (FENE) springs with interaction energy

$$V_{\mathrm{FENE}}(r)=\frac{-K_{\mathrm{FENE}}}{2}R_0^2\ln\left[1-\left(\frac{r}{R_0}\right)^2\right]+V_{\mathrm{WCA}}(r),$$

with $K_{\mathrm{FENE}}=30k_BT/\sigma^2$ being the bond strength and $R_0=1.6\sigma$ the maximum bond length. Third, a bending stiffness is provided by the Kratky-Porod potential

$$V_{\mathrm{BEND}}(\phi_i)=K_{\mathrm{BEND}}\left(1+\cos\phi_i\right),$$

where $\phi_i$ is the angle between beads $i-1$,$i$ and $i+1$, and $K_{\mathrm{BEND}}=4k_BT$ leads to a persistence length comparable to that of chromatin. Finally, additional springs are used to "crumple" the polymer in some regions (*i.e.*, to give it heteromorphic properties). A harmonic spring potential is used, described by

$$V_{\mathrm{HARM}}(r)=K_{\mathrm{HARM}}(r-R_{\mathrm{HARM}})^2,$$

with spring constant $K_{\mathrm{HARM}}=200k_BT/\sigma^2$ and equilibrium bond length $R_{\mathrm{HARM}}=1.1\sigma$. This is applied between next-nearest neighbour beads ($i$ and $i+2$). The regions of the polymer where these springs are included then have a more compacted structure, with regions where they are not included being more open.

Protein complexes are represented by spheres, also of diameter $\sigma$. They interact sterically with each other via the WCA potential as given in Eq. (1). They have a strong attractive interaction with a subset of polymer beads which represent protein binding sites on the chromatin, and a steric (WCA) interaction with all other chromatin beads. The "active" species of proteins additionally have a weak attractive interaction with open chromatin regions (which lack the harmonic "crumple" springs). The attractive interactions are described by a shifted and truncated Lennard-Jones potential

$$V_{\mathrm{LJ/cut}}(r)=\begin{cases}[V_{\mathrm{LJ}}(r)-V_{\mathrm{LJ}}(r_c)] & r\le r_{\mathrm{cut}},\\ 0 & \text{otherwise,}\end{cases}$$

with

$$V_{\mathrm{LJ}}(r)=4\epsilon\left[\left(\frac{\sigma}{r}\right)^{12}-\left(\frac{\sigma}{r}\right)^{6}\right],$$

where $r_{\mathrm{cut}}=1.8\sigma$ is a cut-off distance and $\epsilon$ is the interaction strength. For strong interactions (*i.e.*, with protein binding sites) we use $\epsilon=8k_BT$, while for weak non-specific interactions we use $\epsilon=2k_BT$.

During the simulations proteins switch back and forward between a binding and a non-binding state with at rate $k_{\mathrm{switch}}$ (Brackley *et al.*, 2017). This represents post-translational modifications which alter the protein-DNA binding affinity (*e.g.*, phosphorylation). We set $k_{\mathrm{switch}}=10^{-3}\ \tau^{-1}$, where $\tau$ is the simulation time unit (see below). When in the non-binding state the proteins interact with all chromatin beads via the WCA potential.

#### 1.2 Langevin Dynamics

To perform the simulations we use the LAMMPS molecular dynamics software (Plimpton, 1995) in Langevin dynamics mode. The positions of the polymer beads and proteins evolve according to the Langevin equation

$$m\frac{\mathrm{d}^2\mathbf{r}_i}{\mathrm{d}t^2}=-\boldsymbol{\nabla}_iU-\gamma\frac{\mathrm{d}\mathbf{r}_i}{\mathrm{d}t}+\sqrt{2k_BT\gamma}\boldsymbol{\eta}_i(t), \qquad (2)$$

where $\mathbf{r}_i$ is the position of the $i$th bead. For simplicity we assume all polymer and protein beads have the same mass $m$; $U$ is the total potential energy of the system and $\gamma$ is the friction due to an implicit solvent, again assumed to be the same for all polymer and protein beads. The vector $\boldsymbol{\eta}_i$ is a thermal noise term with components where

$$\langle\eta_{i,\alpha}(t)\rangle=0 \text{ and } \langle\eta_{i,\alpha}(t)\,\eta_{j,\beta}(t')\rangle=\delta_{i,j}\delta_{\alpha,\beta}\delta(t-t'),$$

where $\delta_{i,j}$ is the Kronecker delta and $\delta(t-t')$ is the Dirac delta function.

Equation (2) is solved in LAMMPS using a velocity-Verlet scheme. We use a time step $\mathrm{dt}=0.01\ \tau$, where $\tau$ is the simulation time unit (see below for details of mapping to real times).

#### 1.3 Loop extrusion

The HiP-HoP model includes the loop extrusion mechanism thought to be performed by the cohesin complex (Nishiyama, 2019; Fudenberg *et al.*, 2016; Sanborn *et al.*, 2015). Cohesin complexes are represented by additional spring bonds between polymer beads; the pair of beads to which the bond is applied is then moved outwards, *i.e.*, from $(i,i+3)\to(i-1,i+4)\to(i-2,i+5)$, *etc.*, to extrude a loop. Extruders are added at random positions $i,i+3$ with rate $k_{\mathrm{on}}=0.02\ \tau^{-1}$ and removed with rate $k_{\mathrm{off}}=2.5\times10^{-5}\ \tau^{-1}$; we include 6 extruders per Mbp of simulated chromatin. They are stepped stochastically at rate $k_{\mathrm{ex}}=2\ \mathrm{bp}\tau^{-1}$. The spring potential is given by

$$U_{\mathrm{EXTR}}(r)=U_{\mathrm{WCA}}(r)+K_{\mathrm{EXTR}}(r-r_0)^2,$$

where the bond has strength $K_{\mathrm{EXTR}}=40k_BT/\sigma^2$ and length $r_0=1.5\sigma$. An extruder is halted either when it meets another extruder, or when it meets a CTCF site whose direction is opposite to that of the extrusion. If an extruder is halted on one side, the position at the other side continues to be moved, *i.e.*, it continues to extrude the loop on one side. These parameters were chosen because they were previously shown to generate good predictions of single cell and population average chromatin configurations (Buckle *et al.*, 2018); also the main extrusion control parameter, processivity $k_{\mathrm{ex}}/k_{\mathrm{off}}$, is in line with values previously used in the literature (Goloborodko *et al.*, 2016).

##### 1.4 Simulation setup

For each locus of interest, we simulated a 3 Mbp region of the genome (represented by 3000 polymer beads). In order to improve the efficiency, in each simulation we included 11 copies of the same locus spread across a large polymer of length 40,000 beads (a 600 bead "spacer" region was included between each copy). By combining multiple copies of the locus in this way, we were also able to match the typical chromatin density of the nucleus [this was not possible in Buckle *et al.* (2018), where only a single locus was represented in each simulation].

Initial 40,000 bead configurations were obtained by first arranging the polymer in a mitotic chromosome-like structure similar to that in Rosa and Everaers (2008). This was then relaxed in the absence of protein interactions, but with loop extrusion. This 'equilibration' simulation was run for long enough so that the system had moved away from its initial structure, but retained a looping probability curve consistent with the so-called fractal globule (Mirny, 2011). Crumpling springs were then added according to the input data (see below), before further relaxation.

For the main results, for each locus we performed two independent simulations for a run time of 500,000$\tau$. After discarding the first 100,000$\tau$ to allow the system to reach a steady state, we then extracted a configuration every 2000$\tau$ (obtaining 200 from each simulation). This then gives a total of $11 \times 2 \times 200 = 4400$ configurations for each locus/rearrangement, each of which can be thought of as representing a single cell.

Our simulations have several parameters and the values of these affect the result to a greater or lesser extent. Our previous work (Buckle *et al.*, 2018) suggests that the number (or density) of model proteins, the number of loop extruders, and the speed and unbinding rate of loop extruders have the largest effect on the predicted chromatin interactions. Parameter choices here were guided by those of Buckle *et al.* (2018); it was not always possible to keep the parameters the same, however, due to change in the simulation set-up. Here we simulate several copies of the locus in each simulation, and we use a more realistic chromatin density than in previous work; this affected our choice of the number of proteins, as there is no clear way to map from the previous work (*e.g.*, one could preserve protein concentration or the ratio between proteins and binding sites). Secondly, as detailed below, the mapping between simulation lengths and times and real lengths and times was not fixed in advance in our simulations, but inferred by comparison to data. The higher chromatin density here compared to previous work means that this mapping is different; this meant that difference choices of rate parameters ($k_{\text{ex}}$, $k_{\text{off}}$ *etc.*) were appropriate. We did not perform any parameter optimisation, and so it may be possible to further improve our simulation predictions; however we would not expect this to change any of our conclusions. [Rev3 point 4; Rev 4 point 1]

##### 1.5 Length and time units

In the simulations, lengths are defined relative to the bead diameter $\sigma$, while the simulation time unit follows naturally from the length, mass, and energy scales as $\tau = \sqrt{\sigma^2 m/k_B T}$. Another important time scale is the time it takes a bead to diffuse across its own diameter, the so-called Brownian time, $\tau_{\text{B}} = \sigma^2/D$, where $D$ is the diffusion constant defined through the Einstein relation $D = k_B T/\gamma$. As noted above, for simplicity we set the mass of polymer beads and proteins to be the same, $m = 1$, and we set the friction $\gamma = 2$. With this choice $\tau_{\text{B}} = 2\tau$; the system is therefore over-damped, though the inertial forces are larger than in reality. This approximation is necessary to keep the overall simulation times feasible.

While each polymer bead in the simulations represents 10 kbp of chromatin, it is not necessary to specify a physical size which this occupies. We therefore quote lengths in units of $\sigma$, the bead diameter, throughout. Nevertheless, it is possible to obtain a mapping between simulation and real lengths, *e.g.*, by comparing simulation measurements with 3D FISH measurements. In Buckle *et al.* (2018) this led to a length unit of $\sigma \approx 21.8$ nm.

Simulation time units can be mapped to real units through the Brownian time by calculating the mean square displacement (MSD) as a function of lag time for all polymer beads. This is then compared to experimental data from Hajjoul *et al.* (2013), where MSDs were obtained from various chromatin loci which were fluorescently labelled in live yeast cells. The value for $\tau_{\text{B}}$ which gives the best fit to this data was obtained (using the value of $\sigma$ quoted above for the length unit). We find a mapping where $\tau \approx 5$ ms. This means that a 500,000$\tau$ simulation run maps to approximately 42 minutes.

#### 2 Simulation input data

As detailed in the main text, a number of publicly available data sets are used as an input for the simulations.

First, DNase hypersensitive sites (DHS) are used to identify active protein binding sites. We use the simplifying assumption that there is a single species of active protein (*e.g.*, a complex of transcription factors and RNA polymerase) which binds all DHS with equal affinity. DNase-seq data were obtained from the ENCODE Project for GM12878 cells (The ENCODE Project Consortium, 2012) or the BLUEPRINT Project for U266 and Z-138 cells (Stunnenberg *et al.*, 2016). In both cases DNase "hotspots" (peaks) and "fold enrichment" (FE) data were available. Many hotspots, while having a statistically significant enrichment of reads, nevertheless have very low signal. We therefore further filtered these using a peak threshold and minimum separation criteria based on the FE signal. Since our polymer model has a resolution of 1 kbp we required peaks to have a minimum 1 kbp separation. The centre point of each peak was used to identify the corresponding polymer bead.

Second, a panel of six ChIP-seq data sets for different histone modifications (H3K4me3, H3K4me1, H3K27me3, H3K9me3, H3K27ac and H3K36me3) were used to identify 11 chromatin states via a hidden Markov modelling (HMM) scheme as detailed in Carrillo-de Santa-Pau *et al.* (2017); see also Supplemental Table S1. States for U266 and Z-138 cells were available from the BLUEPRINT Project directly (Stunnenberg *et al.*, 2016). For GM12878 cells we obtained ChIP-seq data from the ENCODE project (The ENCODE Project Consortium, 2012), and generated states using the ChromHMM software [Ernst and Kellis (2012); briefly, ChromHMM segments the genome into 200 bp bins and assigns each to one of the 11 states from the previously learned model from Carrillo-de Santa-Pau *et al.* (2017)]. States associated with H3K27me3 were used to identify binding sites for a polycomb (repressive) model protein (Bantignies and Cavalli, 2011). States associated with H3K9me3 (found in heterochromatin regions) were used to identify binding sites for a second repressive species of model protein [*e.g.*, representing HP1 (Eissenberg and Elgin, 2000)]. States associated with H3K27ac were used to identify the regions of the polymer which have the more open (no "crumpling" springs) structure (Risca *et al.*, 2017). Details of the mapping between states and the simulations are given in Supplemental Table S1. We note that the ChIP-seq data represent histone modifications at both alleles of the locus of interest, and so in this sense the simulations generate structures based on merged information. This is a limitation of the input data rather than the simulation method; if the data could be mapped in an allele-specific way, it would be possible to simulate these separately.

Finally, ChIP-seq data for CTCF obtained from the ENCODE project (The ENCODE Project Consortium, 2012) was used to identify "loop anchors", which halt loop extrusion in a direction depended way. Data was available for the GM12878 cell line; no data was available for the U266 or Z-138 cell lines, so we instead used data from healthy B-cells (CD20+ cells). In all cases we obtained read alignments from ENCODE for "signal" and "input" experiments (in BAM format aligned to the hg19 reference genome), and performed peak calling using the MACS2 software (Zhang *et al.*, 2008). For each called peak we searched the underlying sequence for the CTCF biding motif [matrix MA0139.1 obtained from the JASPAR database (Fornes *et al.*, 2019)] using the FIMO tool (Grant *et al.*, 2011) from the MEME software suit (Bailey *et al.*, 2009), identifying the direction of the motif. Where multiple motifs with different directions were found within a single peak, we used the motif which fit most closely to the consensus (unless motifs with opposite directions scored similarly in comparison to the consensus, in which case the peak was labelled as having both directions). In each simulated copy of the locus a CTCF site is assumed to be bound by a CTCF protein with a probability based on the ChIP-seq peak height (*i.e.*, this varied between copies, accounting for cell-to-cell variation in CTCF binding).

Details of the source of each data set, including paper references and accession numbers where appropriate, are given in Supplemental Table S3

#### 3 Simulation Measurements

As noted above, for each locus/rearrangement we generated 4400 independent 3D configurations. These provide the positions in simulation space of each 1 kbp chromatin bead which makes up the 3 Mbp (3000 bead) region of interest.

To obtain a simulated Hi-C interaction map, the procedure was as follows. Two chromatin beads were selected at random, and their separation calculated; this was then counted as a contact with probability $P_{\text{contact}} = \exp(-r/r_c)$, where $r$ is the separation and $r_c$ is a threshold length scale which we set to $2\sigma$. This mimics a Hi-C experiment where the likelihood of two loci being joined during fixation and subsequently becoming ligated decreases with 3D separation. This was repeated $N(N-1)/2$ times for each configuration so that each possible pair of $N$ beads had the chance to be selected. This entire process was then repeated enough times such that the number of contacts found was of the same order of magnitude as the number of reads in an experimental Hi-C map of the same size (*i.e.*, we approximately matched the read depth). Reads were then organised into 3 kbp bins and contact maps plotted using the HiCExplorer software (Ramírez *et al.*, 2018).

To obtain simulated 4C-like profiles we first identified a set of 'bait' (or 'target') beads, before following a similar procedure as for the Hi-C maps. For each bait, in each simulated configuration we picked a bead at random, and calculated the separation between it and the bait. We counted it as a contact with probability $P_{\text{contact}}$ as above; this was done $N$ times so that each bead had the chance to be selected. A profile was then generated at 1 kbp resolution.

To measure the 3D size of a given region of the genome, we computed the radius of gyration, $R_g$, of the beads which made up that region. This is defined by

$$R_g^2 = \frac{1}{M}\sum_i (\mathbf{r}_i - \mathbf{r}_{\text{CoM}})^2,$$

where $\mathbf{r}_i$ is the position of bead $i$, the sum runs over the $M$ beads within the region of interest, and $\mathbf{r}_{\text{CoM}}$ is the centre of mass of these beads, as given by

$$\mathbf{r}_{\text{CoM}} = \frac{1}{M}\sum_i \mathbf{r}_i.$$

A more accurate measure of the 3D size of a region which takes some account of the shape was used in Fig. 6G in the main text. A gyration tensor can be defined, where the $\alpha,\beta$ element is given by

$$G_{\alpha\beta} = \frac{1}{N}\sum_{i=1}^{M} r_\alpha^{(i)} r_\beta^{(i)},$$

where $r_\alpha^{(i)}$ is the $\alpha$th component of the vector $(\mathbf{r}_i - \mathbf{r}_{\text{CoM}})$, and the sum runs over the $M$ beads within the region (gene) of interest. The sum of the eigenvalues of this matrix is equal to $R_g^2$, and the eigenvalues themselves can be thought of as the squares of the lengths of the principal axes of an ellipsoid. The volume of this ellipsoid gives a measure of the volume the region occupies in 3D space.

#### 4 Chromosome-conformation-capture data for comparison with simulations

Hi-C data for GM12878 (Fig. 2Ai and Supplemental Figs. S1 and S2) was obtained from Rao *et al.* (2014). We used the processed data available from that reference using the "square root vanilla coverage" normalisation as detailed therein. Hi-C data (raw unprocessed paired-end reads) for U266 cells (Fig. 4C and Supplemental Fig. S4) was obtained from Wu *et al.* (2017). For comparison with our simulations we generated a U266 reference genome based on hg19, which included the chromosome 14 *IGH* enhancer region inserted near *CCND1* on chromosome 11 (see section below for details of how the rearrangement was mapped). In other words, we generated an *in silico* rearranged reference genome. Though we expect U266 cells to harbour additional rearrangements and point mutations, including this single insert is sufficient for comparison with our simulations. To avoid having reads map to multiple locations we did not include chromosome 14 in our reference. We note that the Hi-C data set will include interaction reads from both copies of these chromosomes (those with the rearrangement and those without). We proceeded to map reads and generate Hi-C interaction maps using the HiC-Pro software (Servant *et al.*, 2015), which employs Bowtie 2 (Langmead and Salzberg, 2012) for read mapping, and the ice matrix normalisation method (Imakaev *et al.*, 2012). The Wu *et al.* (2017) paper provides data from experiments using HindIII and MBoI restriction enzymes, which we analysed separately, and then combined. The GM12878 dataset has a high read depth (4.9 billion reads), allowing interaction maps to be generated with a resolution of 10 kbp bins; the U266 data had a lower read depth (228 million reads), but a lower 20 kbp resolution enabled the same downstream analysis to be performed.

We note that, importantly, Hi-C data was not used as an input to the simulations, only as a comparison. The simulations therefore predict 3D structure based only on 1D genomic information. To compare domain patterns between the simulations and experimental Hi-C data, we used the Hi-C Explorer software (Wolff *et al.*, 2020) to call TAD boundaries. As with most boundary algorithms, calls depend strongly on bin size and read depth/noise, and several parameters need to be tuned. For an unbiased comparison, we tuned parameters to obtain TAD calls which closely matched the visible domains on the experimental interaction maps, and then used the same parameters for the simulated maps (with the latter using the same bin size as the former). Supplemental Fig. S2B shows TAD calls for the GM12878 cells overlaid on an interaction map. From the experimental data there were 6 boundary calls; from the simulation data there were 5 boundary calls which matched those from the experiment to with 2 bins. The simulation showed two additional boundaries which did not exactly match the experiment (green arrows in Supplemental Fig. S2B); while this suggests that the simulations performed less well in this region, a visual comparison of the maps show there is still reasonable agreement, and that the discrepancy could instead be a result of poorer boundary calling in that region. TAD calls for the U266 cells are shown in Supplemental Fig. S4B; again, there were 6 boundaries called from the experimental data. The simulations also showed 6 boundaries, with 5 matching to within 2 bins. The boundary which does not match (cyan arrow in Supplemental Fig. S4B) is next to an unmappable DNA region, which has likely caused the boundary calling to fail; a more likely position for the boundary is shown by the purple arrow in the figure, and this does match that found in the simulations. We also note a 'stripe' of enriched interactions between *CCND1* and the region to its left in both simulations and data (green ellipses in Supplemental Fig. S4B). Together this suggests that the *CCND1* gene does indeed fall within the TAD which encompasses the insert region, as predicted by the simulations.

For a more holistic comparison between the experimental and simulation Hi-C maps, we considered correlations between experimental and simulated interaction maps. There is a strong drop off in Hi-C interaction with genomic separation (see Supplemental Fig. S2C), so to obtain a measure which is independent of this, we considered the interactions between all points at a *given* genomic separation (equivalent to taking a line across the interaction map parallel to the diagonal, Supplemental Fig. S2E). We calculated the Pearson correlation coefficient between this signal from experiments and simulations, and then plot as a function of genomic separation. Supplemental Figure S2E shows this for the GM12878 cell line; we found statistically significant correlation coefficients of around 0.5 over a wide range of genomic separations. To put this into context we also consider two 'control' cases. First, we generated a 'shuffled' interaction map by randomly swapping interaction intensities in a way which preserves the dependence on genomic separation (the resulting map is devoid of TAD or other patterns, as shown in Supplemental Fig. S2D). Second, we generated a 'shifted' map, where all interaction were shifted to the right by 250 kbp (the resulting maps has TAD boundaries and interaction spots in different locations, Supplemental Fig. S2D). Supplemental Figure S2E shows that there is very little correlation between these controls and the experimental map (Pearson correlation coefficients are close to zero, and the correlation is not significant, $p > 0.01$). Similar results were obtained for the U266 cell line (Supplemental Fig. S4C), though here the lower read depth/noisier data led to lower correlation values.

We note that while the correlation measure detailed above is not affected by the strong drop off in interactions as genomic separation increases (Supplemental Fig. S2C), it is highly sensitive to noise (particularly at large genomic separations). As an alternative, we also considered the 'directionality index' (DI), which is often used to quantify domain insulation [e.g, in Dixon *et al.* (2012)]. The DI at bin $i$ in a Hi-C map is defined as

$$\mathrm{DI}_i = \left( \sum_{j=i+1}^{i+w} H_{ij} \right) - \left( \sum_{j=i-w}^{i-1} H_{ij} \right),$$

where $H_{ij}$ is the Hi-C count of interactions between bins $i$ and $j$, and $w$ is a window length. In other words, the DI is the difference between the level of interactions to the right and to the left of a point within a distance of $w$ bins (see also Supplemental Fig. S2F). For a given value of $w$ we can find a DI profile for both experiments and simulations, and then calculate the Pearson correlation coefficient. Supplemental Fig. S2F shows a plot of the correlation coefficient as a function of window length. For small window lengths, this measure is also sensitive to noise; however, as $w$ increases it becomes more robust (the measure effectively averages over the window). As expected, the DI correlation increases with window length, reaching values of around 0.7-0.75; this drops off again as the window size gets larger, due to the finite size of the simulated region. For all points in Supplemental Fig. S2F, the correlation is statistically significant ($p < 0.01$). Also show is the correlation for the shuffled and shifted maps as detailed above; in these cases the values are close to zero, and the correlation is not significant ($p > 0.01$). Again, similar results were observed for the U266 cell line (Supplemental Fig. S4C); while the correlation is smaller due to the lower read depth, values still reached 0.5.

ChIA-PET data for Rad21 (a cohesin subunit) in GM12878 cells was obtained from Heidari *et al.* (2014), and Hi-ChIP data for H3K27ac in GM12878 cells was obtained from Bhattacharyya *et al.* (2019). Both of these methods combine mapping of chromatin interactions with an immunoprecipitation step, thus providing information on interactions between regions which bind a specific protein or possess a specific histone modification. They therefore provide a more targeted method than Hi-C, and so give higher resolution. In Supplemental Fig. S2G we show interactions from these data sets as 'arcs' alongside a section of a simulated interaction map. As noted in the main text, Hi-C maps tend to show rather uniform interactions within TADs. Our simulations often show more details, with dark spots indicating interactions between, *e.g.*, promoters and enhancers. Supplemental Fig. S2G shows that many of the predicted interactions for the promoters of *CCND1* and nearby *LTO1* are indeed present in the more targeted data sets. Calling significant interactions from 3C based data typically requires building a genome wide background model, which is not possible from simulations which include only a 3 Mbp region; a direct comparison between the simulations and Hi-ChIP/ChIA-PET arcs is therefore not possible. Nevertheless, we can show quantitatively that the simulations do predict enrichment of interactions at these arcs. To do this, first we find a normalised interaction score for all pairs of Hi-C bins in our simulation map by dividing by the mean interaction level for the given genomic separation (*i.e.*, we scale out the strong dependence shown in Supplemental Fig. S2C). We can then compare the distribution of normalised interaction scores for the whole simulated region, with the subset of interactions at the arcs from the ChIA-PET dataset. Supplemental Figure S2G top right shows that there is a clear enrichment of interactions for this set of arcs in the simulations (a Kolmogorov-Smirnov test rejects the null hypothesis that the distributions are the same, $p < 10^{-12}$). We also considered two 'negative controls': first, we randomly rewired all of the arcs from the Hi-ChIP data set, preserving the arc bases; and second, we randomly repositioned all of the arcs within the 3 Mbp region, preserving the arc lengths. Supplemental Figure S2G bottom right shows that the two sets of control arcs do not show enrichment of interactions in the simulations (in both cases, a Kolmogorov-Smirnov test rejects the null hypothesis that the distributions are the same, $p < 10^{-5}$).

#### 5 Cell lines

Here we provide some additional details on the cancer cell lines. The Z-138 cell line was derived from a patient initially diagnosed with chronic lymphocytic leukaemia. Around two years later, the disease transformed into an aggressive mature B-cell malignancy being classified as mantle cell lymphoma with blastoid transformation. Importantly for this study, the cells harbour a reciprocal translocation, t(11;14)(q13;q32)/*IGH-CCND1* resulting in overexpression of cyclin D1. The breakpoint for this translocation within the *IGH* locus, is located in the J gene region close to the *IGH* E$\mu$ super-enhancer, suggesting the translocation arose in an immature B-cell undergoing V(D)J rearrangement.

The U266 cell line was derived from a patient with refractory and terminal myeloma. On the background of a complex karyotype, this cell line harbours an insertion of chromosome 14q32 material into chromosome 11q13. This insertion results in the relocation of the E$\alpha$1 *IGH* super-enhancer into close proximity to the *CCND1* gene at 11q13. Again, this insertion event leads to overexpression of cyclin D1. The breakpoints within the *IGH* locus reside in the constant region, suggesting this rearrangement occurred when the cell was undergoing class switch recombination to result in the production of IgE-secreting plasma cells.

#### 6 Gene expression measurements

To understand how the level of expression of *CCND1* in GM12878 cells compares with expression levels in general, we obtained RNA-seq data from the ENCODE project [assession ENCSR889TRN; The ENCODE Project Consortium (2012)]. Specifically, we used the ENCODE gene expression quantifications, which provide a 'fragments per kilo-base of transcript per million mapped reads' (FPKM) measure. As a reference point we considered the set of standard housekeeping genes, previously identified by Eisenberg and Levanon (2013), namely *C1orf43*, *CHMP2A*, *EMC7*, *GPI*, *PSMB2*, *PSMB4*, *RAB7A*, *REEP5*, *SNRPD3*, *VCP* and *VPS29*. We found that in GM12878 cells, *CCND1* has an FPKM which is 105.3 times smaller than the average of these housekeeping genes.

To further quantify gene expression across the three cell lines, we performed qPCR as detailed in the Methods section in the main text. The oligonucleotide sequences are shown in Table S2. Some additional results are shown in Supplemental Fig. S3; for each cell line we show the relative expression level for each gene, using that of *TPCN2* as a reference [Supplemental Figs. S3Ai, Bi and Ci].

Our simulations predict the configuration of the gene locus in 3D, providing a set of structures which each represent a single cell. Exactly how 3D structure affects the expression of a given gene remains the subject of current research (Kempfer and Pombo, 2020). For some genes it has been shown that physical interaction between a promoter and an enhancer, mediated by proteins bound at these elements, is required to activate transcription (Schoenfelder and Fraser, 2019). Based on this we hypothesised that our simulated structures could also be used to predict gene expression level.

In previous work using a simpler polymer model for chromatin (Brackley *et al.*, 2020) we measured the fraction of time during a simulation which a promoter region is bound by an active protein (in the simulations the active proteins represent a general complex of transcriptional activators, transcription factors, and/or polymerase), and found that this correlates with gene expression as measured by GRO-seq experiments. This is a very simple predictor measure, since it does not take into account enhancer "strength" or compatibility, nor any further biochemical regulatory mechanisms (*e.g.*, transcription factor specificity). We therefore cannot expect a clear mapping between the predictor and actual transcription level, only a correlation. We performed similar measurements for the present simulations (Supplemental Fig. S3Di); a protein was defined as being bound to a promoter if the separation between the protein bead and polymer bead was less than $2.25\sigma$. This measure did not give particularly good predictions of relative expression levels within the same cell line, however the predicted difference between cell lines was better (Supplemental Fig. S3Ei and ii, left panels). This is not surprising, given the limitations discussed above. We also considered a second predictor for transcription, instead asking how often the promoter of a gene interacts with another chromatin region which has an enhancer associated state (an "interaction" was again defined as when the two beads were closer together than $2.25\sigma$, and we also specified that the enhancer must be at least 4 kbp away from the promoter). Such interactions/loops tend to arise due to proteins binding at both elements to form bridges (and clusters of proteins), though they can also be driven by the loop extrusion mechanism. This measure gave even better predictions of the experimentally measured differences in expression between the cell types (Supplemental Fig. S3Ei and ii, right panels and Fig. 6C in the main text). We note though that the same limitations as discussed above apply.

#### 7 Fluorescence *in situ* hybridisation microscopy

As detailed in the Methods section in the main text, we performed fluorescence *in situ* hybridisation microscopy experiments on the three cells lines. More specifically, fosmid clones found to cover TPCN2 (G248P87917D11/WI2-1721H21), MYEOV (G248P87014D2/WI2-2222G4) and CCND1 (G248P86668E2/WI2-2191I3) loci were obtained from BACPAC resources (California, USA). Single bacterial colonies were cultured in the presence of chloramphenicol and DNA extracted using the GeneJET Plasmid Maxiprep Kit (Thermo Scientific, Massachusetts, USA). 1$\mu$g of DNA was labelled by nick translation using the Enzo Nick translation

DNA labelling system 2.0 (Enzo Life Sciences, Exeter, UK) and fluorophores SEEBRIGHT Green 496 dUTP, SEEBRIGHT Red 598 dUTP or SEEBRIGHT Gold 525 dUTP. The resulting fluorescently labelled probes were mixed 1:1 with hybridisation buffer (Cytocell, Cambridge, UK ) and denatured at 75°C for five minutes followed by hybridisation at 37°C overnight. Slides were washed for two-minutes in 0.02% SSC with 0.003% NP40 at 72°C followed by two minute room temperature incubation in 0.1% SSC. Slides were mounted with 10$\mu$l DAPI (Vector laboratories, California, USA).

Images were acquired on a Leica SP8 point scanning confocal microscope with white light super continuum lasers (tunable from 470 nm to 670 nm and a fixed 405 nm Argon laser using a HC PL APO CS2 63×/1.40 OIL lens. Images were acquired as Z-stack for measurement and analysis of 3D volumes. Voxel size for all images was set at 43×43×130 nm $(x, y, z)$ to allow deconvolution and consistent particle analysis (where a 'particle' is a contiguous region within which a given probe fluorescence intensity was above a threshold). All the other settings were also kept consistent between experiments, specifically: DAPI (for nuclear counterstain), Ex 405 nm – Em 410/480nm, Seebright-green, Ex 496 nm – Em 502/525 nm, Seebright-gold, Ex 525 nm – Ex 531/592 nm, Seebright-red-598-dutp, Ex 596 nm – Em 606/781 nm. Images were deconvolved using Huygens 20.04 software (www.svi.nl). Particle analysis was performed using Huygens 20.04 on deconvolved images as recently shown (Prendergast *et al.*, 2020). Specifically, we measured 'Small Particle Geometry', obtaining the volume in voxels.

We found that these large fosmid probes were difficult to work with in these cell lines, and this limited the number of imaged cells available. Often, in a given cell we would observe more than the two expected particles (one per chromosome copy). This was likely due to hybridisation problems (the probe not hybridising uniformly, so the region appears as multiple distinct particles), or because different cells were at different stages of the cell cycle (particles from different sister chromatids being visible in G2, or different chromosomes being replicated to different extents in S-phase). It was difficult to find cells where there were exactly two particles for each probe. In cases when this was the case, it was not always clear which particles were on the same chromosome copy. Nevertheless, we were able to obtain volume measurements for at least 20 cells (40 particles) for each probe in each cell line, as shown in Fig. 6F and Supplemental Figs. S10B, E and D. The differences between probe volumes in different cell lines showed similar trends to those predicted in simulations (namely, volumes for *CCND1* and *MYEOV* probes tended to be larger in the cancer cell lines, Supplemental Figs. S10A, D and G). However, with these numbers of cells, there was insufficient statistical power to reject the null hypothesis that the volumes were drawn from the same distribution (via a two-sample Kolmogorov-Smirnov test).

Another feature of the FISH measurements, is that (with the exception of Z-138 cells) it is not possible to distinguish particles from the rearranged or non-rearranged chromosomes. Under the assumption that one of the *CCND1* alleles will behave as in a healthy cell, we can construct a more realistic prediction of the volume trends by combining simulated measurements from the cancer and healthy cell lines. These predictions are shown in Supplemental Fig. S10C, F and I. We have also 'downsampled' the data so that the number of measurements matches that of the experimental data; i.e., we pick a random subset of measurements from a given simulation. For example, for the green box in Supplemental Fig. S10C we randomly picked 23 measurements from the U266 simulations and 23 measurements from the GM12878 measurements, to match the 46 measurements from the U266 experiment (where half of the particles are from non-rearranged chromosomes). A similar downsampling is done for the other two cell lines (in GM12878, all measurements are taken from the GM12878 simulations). After this procedure, we observe the same general trends as with the full simulation data set, but now there is not sufficient statistical power to reject the null hypothesis. In this respect, the trends observed in our simulations are consistent with the FISH data (as expected with a simple model, we obtain qualitative rather than quantitative spatial predictions). [rev1 point2;]

#### 8 Break-point mapping in U266 and Z-138 cells

Chromosomal break-points of t(11;14) in U266 (Mikulasova *et al.*, 2020b) and Z-138 cell lines were detected using paired-end read targeted sequencing of the DNA involving extensive coverage of the *IGHC*, *IGHJ*, and *IGHD* loci (~300 kbp) (Mikulasova *et al.*, 2020a). FASTQ files were aligned to the human genome assembly GRCh37 using BWA-MEM (Li, 2013). Chromosomal rearrangements were called using Manta (Chen *et al.*, 2015) with default settings. Detected break-points were manually inspected in the Integrative Genomics Viewer [Thorvaldsdóttir *et al.* (2013); Broad Institute, Cambridge, MA, USA]. The sequencing data are available at the Sequence Read Archive (SRA), National Center for Biotechnology Information (NCBI, Bethesda, MD, USA) under the BioProject accession number PRJNA635269.

### Supplemental Tables

| State | Description | Use in simulations |
|---|---|---|
| 1 | Transcription low signal H3K32me3 | – |
| 2 | Transcription high signal H3K32me3 | – |
| 3 | Heterochromatin high signal H3K9me3 | heterochromatin protein binding |
| 4 | Low signal | – |
| 5 | Heterochromatin high signal H3K27me3 | polycomb protein binding |
| 6 | Heterochromatin low signal H3K27me3 | – |
| 7 | Repressed polycomb promoter high H3K4me3, H3K4me1, H3K27me3 | polycomb protein binding |
| 8 | Enhancer high signal H3K4me1 | 'open' chromatin |
| 9 | Active enhancer high signal H3K4me1, H3K27ac | 'open' chromatin |
| 10 | Distal active promoter high signal H3K4me3, H3K4me1, H3K27ac | 'open' chromatin |
| 11 | Active TSS high signal H3K4me3, H3K27ac | 'open' chromatin |

**Supplemental Table S1. Chromatin states.** Table showing chromatin states obtained from the HMM analysis, and how they are used in simulations

| Gene | Sequence | |
|---|---|---|
| ORAOV1 | ATCGTGATGGCGGATGAGAG | CTTCCCTCCATCACACCCAA |
| MYEOV | GTTCTTGGTGGGATCTGAGC | GACACACCACGGAGACAATG |
| TPCN2 | ACTTACCGCAGCATCCAAGT | TTCGATGAAGACCACAGCCT |
| CCND1 | CTGTGCTGCGAAGTGGAAAC | TCACATCTCCAGCATCCAGG |
| GAPDH | GGAAGATGGTGATGGGATTT | GGATTTGGTCGTATTGGG |

**Supplemental Table S2. Oligonucleotide sequences used in qPCR experiments.**

| Cell line | Experiment | Source | Accession | Reference |
|---|---|---|---|---|
| GM12878 | ChIP-seq CTCF | ENCODE | ENCSR000AKB | The ENCODE Project Consortium (2012) |
| GM12878 | DNase-seq | ENCODE | ENCSR000EMT | The ENCODE Project Consortium (2012) |
| GM12878 | ChIP-seq H3K4me3 | ENCODE | ENCFF788INU | The ENCODE Project Consortium (2012) |
| GM12878 | ChIP-seq H3K4me1 | ENCODE | ENCFF753GZX | The ENCODE Project Consortium (2012) |
| GM12878 | ChIP-seq H3K27ac | ENCODE | ENCFF197QHX | The ENCODE Project Consortium (2012) |
| GM12878 | ChIP-seq H3K9me3 | ENCODE | ENCFF331ODM | The ENCODE Project Consortium (2012) |
| GM12878 | ChIP-seq H3K36me3 | ENCODE | ENCFF750QYL | The ENCODE Project Consortium (2012) |
| GM12878 | ChIP-seq H3K27me3 | ENCODE | ENCFF622QTG | The ENCODE Project Consortium (2012) |
| GM12878 | Hi-C | GEO | GSE63525 | Rao *et al.* (2014) |
| whole B-cell | ChIP-seq CTCF | ENCODE | ENCSR000AUV | The ENCODE Project Consortium (2012) |
| U266 | DNase-seq | BLUEPRINT | – | Stunnenberg *et al.* (2016) |
| U266 | Chromatin states | BLUEPRINT | – | Stunnenberg *et al.* (2016) |
| U266 | Break-point mapping | Sequence Read Archive (SRA), National Center for Biotechnology Information (NCBI, Bethesda, MD, USA) | PRJNA635269 | Mikulasova *et al.* (2020b) |
| U266 | Hi-C | GEO | GSE87585 | Wu *et al.* (2017) |
| Z-138 | DNase-seq | BLUEPRINT | – | Stunnenberg *et al.* (2016) |
| Z-138 | Chromatin states | BLUEPRINT | – | Stunnenberg *et al.* (2016) |

**Supplemental Table S3. Data sources.** Table showing source and accession numbers for all publicly available data used in this work. Reference numbers are those given in this Supplemental Material document.

| Data | Availability | url | Accession |
|---|---|---|---|
| Simulation data | Edinburgh DataShare repository | https://datashare.ed.ac.uk | pending |
| Z-138 sequencing data | Sequence Read Archive (SRA), National Center for Biotechnology Information (NCBI, Bethesda, MD, USA) | https://www.ncbi.nlm.nih.gov/sra | pending |
| qPCR data | Edinburgh DataShare repository | https://datashare.ed.ac.uk | pending |
| FISH measurement data | Edinburgh DataShare repository | https://datashare.ed.ac.uk | pending |

**Supplemental Table S4. Data availability.** Table showing availability of all new data generated in this work.

### Supplemental Figures

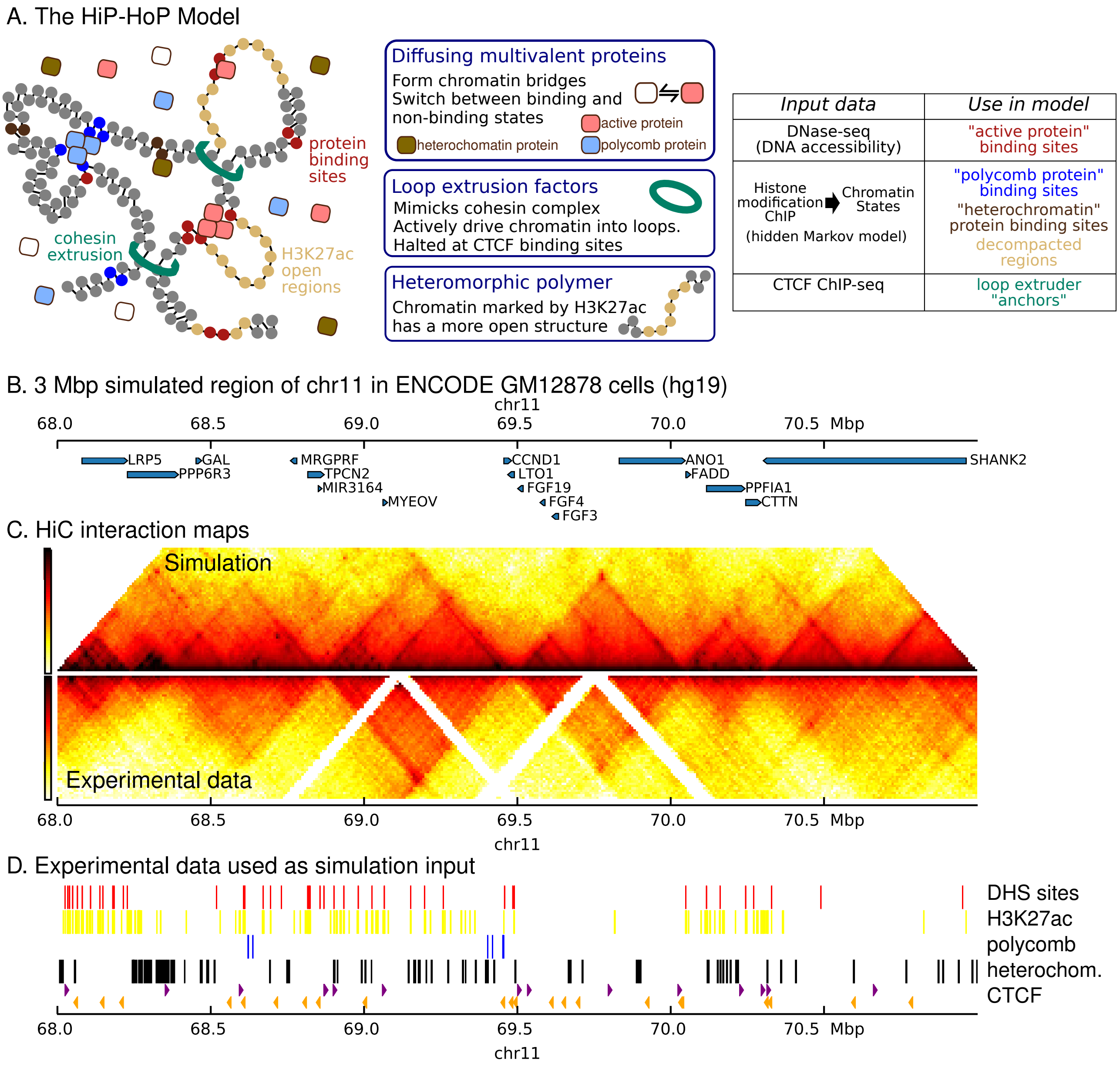

**Supplemental Figure S1. Model schematic with input data and output (simulated Hi-C). A.** Schematic of the HiP-HoP model (left), showing the three included mechanisms which drive 3D structure (middle), with a table describing the simulation input data (right). Three species of diffusing proteins interact with a bead-and-spring polymer representing a chromatin region; binding sites are identified using chromatin states and DNase hypersensitive sites. Chromatin states are also used to identify regions which have a more open decompacted chromatin fibre structure. The loop extrusion mechanism is incorporated, with CTCF ChIP-seq data used to identify anchor sites. **B.** Map showing the gene positions within the 3 Mbp region of human chromosome 11 which was simulated. **C.** Hi-C interaction maps for GM12878 cells from simulations (top) and experimental data [bottom; obtained from Rao *et al*. (2014)]. **D.** Data from the GM12878 cell line used as an input to simulations. Red blocks show DNase hypersensitive sites (DNase-seq data obtained from the ENCODE project (The ENCODE Project Consortium, 2012)). Yellow, blue and black blocks show regions identified as being associated with H3K27ac, polycomb (H3K27me3) and heterochromatin chromatin states from a hidden Markov model (HMM) as detailed in Supplemental Methods. Arrowheads indicate the position and direction of CTCF sites [purple and orange arrowheads indicate left and right facing sites respectively; ChIP-seq data for CTCF obtained from the ENCODE project, The ENCODE Project Consortium (2012), and the direction was determined as detailed in Supplemental Methods].

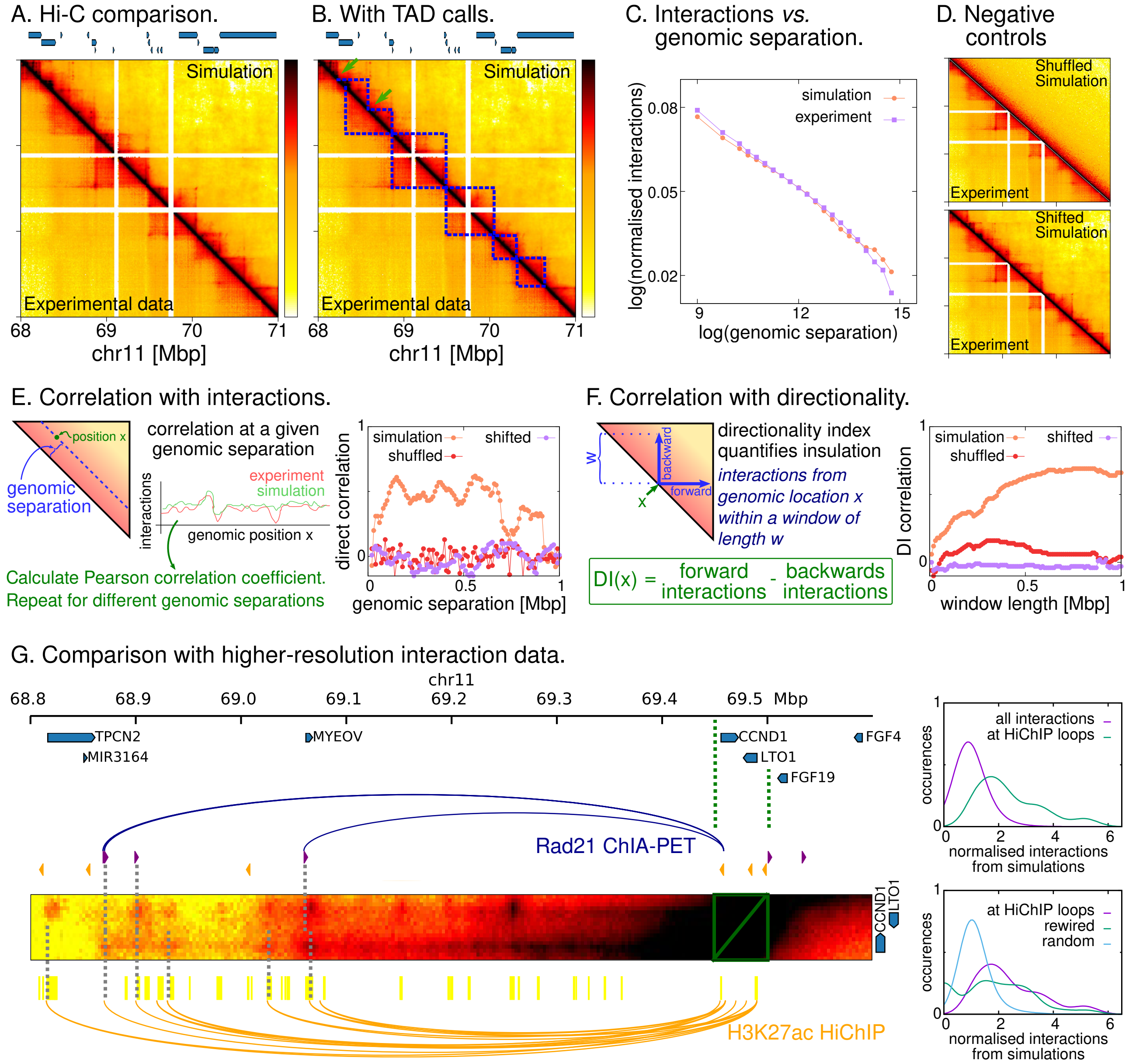

**Supplemental Figure S2. Simulations of *CCND1* in GM12878 cells: quantitative comparison with experiments. A.** Simulated Hi-C is shown alongside the experimental data obtained from Rao *et al*. (2014) for the whole of the 3 Mbp region which was simulated. White gaps in the maps appear at unmappable (repetitive sequence) DNA regions; for ease of comparison, where no data are available from the experiments, the same regions have been masked for simulations. Positions of genes are shown above the plot. **B.** A similar Hi-C map is shown, with overlaid blue dashed lines showing TADs called from the experimental (bottom left) and simulation (top right) maps as detailed in the Supplemental Methods. Green arrows show the position of TAD boundary calls which do not match between simulations and data. **C.** Plot showing interactions as a function of genomic separation on a log-log scale. Separations are binned on a logarithmic scale, and each point represents an average over interactions between all pairs of locations with separations falling within the bin. Simulation and experimental data are show; in the latter case, only interactions within the 3 Mbp regions which was simulated are included. The interaction level is normalised such that the area under each curve sums to unity. **D.** As detailed in the Supplemental Methods, to put quantitative comparisons into context, we consider two 'negative control' cases. Top: the simulation data are shuffled randomly, but preserving the decay with genomic separation. Bottom: the simulation map is shifted laterally by 250 kbp, and periodically wrapped. This results in a map which is different from the original simulation map, but still shows domains and interaction peaks. **E.** Left: schematic showing how an interaction profile can be obtained for all pairs of point at a given genomic separation. The Pearson correlation coefficient between simulated and experimental profiles can be obtained, again for a given genomic separation. Right: the correlation coefficient is plotted as a function of genomic separation. For over 70% of the points, the shown correlation is statistically significant ($p < 0.01$). For context, correlations between the experiment and the shuffled and shifted maps are also shown. These show values close to zero, and are statistically significant for fewer than 5% of the points.

*Caption continued on next page.*

*Continued caption to* **Supplemental Figure S2. Simulations of *CCND1* in GM12878 cells: input data and quantitative comparison with experiments.** **F.** Left: the directionality index (DI) measures insulation as the difference between forward and backward interactions out to a window length $w$ (see Supplemental Methods for details). A profile of DI as a function of position, $x$, can be obtained and the Pearson correlation coefficient between simulated and experimental profiles calculated. Right: the correlation is shown as a function of window length. For all points the correlation is statistically significant. Also shown are correlations for the shifted and shuffled maps; again these have values close to zero, and in this case none are statistically significant. **G.** Comparison between interactions in simulations and higher resolution experimental data. Arcs show ChIA-PET data from cohesin sub-unit Rad21 [blue, obtained from Heidari *et al*. (2014)] and Hi-ChIP data using an antibody for H3K27ac [orange, obtained from Bhattacharyya *et al*. (2019)]. Arcs are shown for interactions with one end within the region bounded by the green dashed lines. The heat map shows a strip from the simulated Hi-C map, with the green box around the diagonal; this therefore shows interactions with *CCND1* and *LTO1*. Grey dashed lines pick out some interactions from the data which are correctly predicted by the simulations. Top right: plot showing how the distribution of interaction signal strengths predicted from simulations for Hi-ChIP loops compares to all interactions. Here interaction strength is normalised so as to remove the dependence on genomic separation; see Supplemental Methods for full details. Bottom right: plot showing how the distribution of interaction signal strengths predicted from simulations for Hi-ChIP loops compares to two 'negative control' cases. First, if loops are rewired (i.e., arcs are randomised, but the same arc bases are used). Second, if arcs are randomly position, but the same arc lengths are used. Full details are given in Supplemental Methods Section 4.

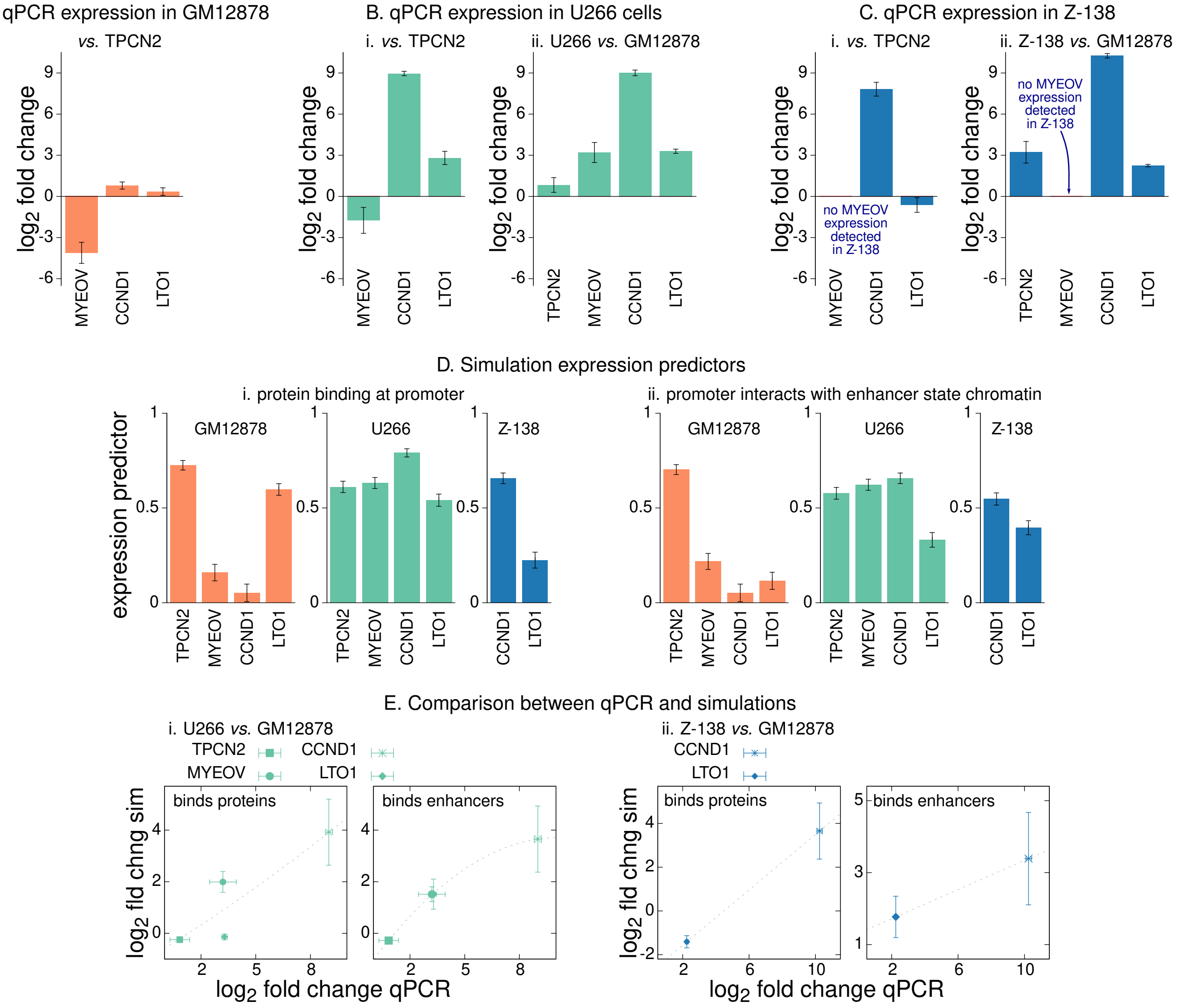

**Supplemental Figure S3. Measuring expression of genes in the *CCND1*-TAD. A.** Bar plot showing expression levels of *MYEOV*, *CCND1*, and *LTO1* using *TPCN2* expression as a reference, obtained from qPCR experiments in GM12878 cells. Values are expressed as the $\log_2$ fold-change and are an average over three replicate experiments; error bars show the standard error in the mean. **B.** Panel i shows a similar plot of expression level using *TPCN2* as a reference for U266 cells. Panel ii shows the expression of all four genes in U266, relative to the expression in GM12878 cells. **C.** Panel i shows the expression in Z-138, again using *TPCN2* as a reference, while in panel ii the expression level in GM12878 is used as a reference. **D.** Plots showing the values of predictors of expression obtained from the simulations as detailed in the main text and Supplemental Methods. Panel i shows the fraction of simulated configurations where the gene promoter was bound by an 'active protein'. Panel ii. shows the fraction of simulated configurations where the promoter was interacting with a distal (*i.e.*, at least 4 kbp away) chromatin region which has an enhancer associated state. Error bars show the standard error for the fraction of 440 configurations. **E.** Plots comparing the *change* in expression compared to GM12878 as predicted from the simulations and as determined experimentally via qPCR. Panels i and ii show data for U266 and Z-138 cells respectively. Left hand plots show the fold change as predicted by proteins binding at the promoter, while right hand plots the fold change predicted by interaction with distal enhancer state chromatin. We do not necessarily expect a linear relationship between simulation and experimental values, but a good correlation is implied if the points lie close to a monotonically increasing curve (see main text and Supplemental Methods; grey dotted lines show such curves as a guide to the eye).

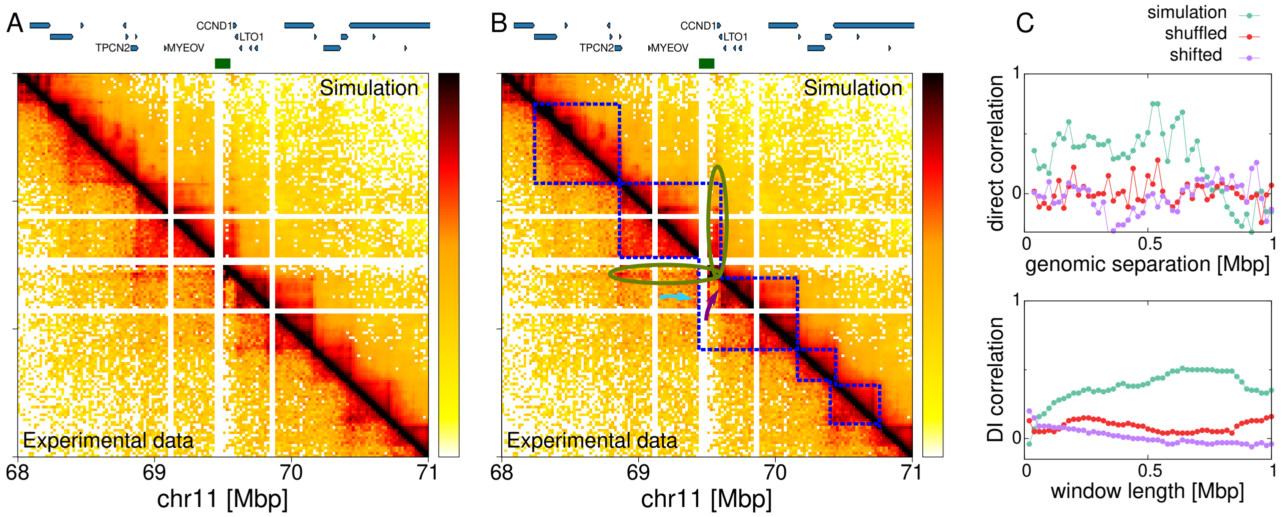

**Supplemental Figure S4. Quantitative comparison between simulated and experimental Hi-C maps for the U266 cell line. A.** Hi-C interaction map for the 3 Mbp region around *CCND1* which was simulated; data are shown in 20 kbp bins (the significantly lower read depth compared to the GM12878 case necessitates this lower resolution). Top right is obtained from simulations; bottom left are Hi-C data for U266 cells obtained from Wu *et al.* (2017). Reads were mapped to an *in silico* re-arranged U266 genome as detailed in the Supplemental Methods Section 4. White gaps in the maps appear at unmappable (repetitive sequence) DNA regions; for ease of comparison, where no data are available from the experiments, the same regions have been masked for simulations. Positions of genes are shown above the plot, and the green block above the plot indicates the position of the insert region. **B.** A similar Hi-C map is shown, with several features overlaid. Blue dashed lines show TADs called from the experimental (bottom left) and simulation (top right) maps as detailed in the Supplemental Methods. In the experiments, the position of one TAD boundary has been called at the edge of an unmappable region where not data are available (cyan arrow). Due to these gaps in the data, a nearby visible boundary (purple arrow) was not called. Green ellipses show an enrichment of interactions between *CCND1* with the TAD to its left, strongly suggesting that this gene sits within this TAD. **C.** Plots showing quantitative comparisons between the simulation and experimental Hi-C maps (see Supplemental Fig. S2E-F for schematic). Top: the interaction signal for all pairs of points at a given genomic separation was obtained, and the Pearson correlation coefficient comparing the simulation and experiment calculated. This is plotted as a function of genomic separation (green points; for 38% of points the correlation is statistically significant, $p < 0.01$). Also shown is the same measure for a shuffled and a shifted contact map (see Supplemental Methods and Supplemental Fig. S2D for details; the values of the correlation coefficient are close to zero, and for only 8% of these points was the correlation significant). Bottom: the directionality index for a given window width was obtained, and the Pearson correlation coefficient comparing the simulation and experiment calculated. This is plotted as a function of window width (green points; for 86% of points the correlation is statistically significant, $p < 0.01$). Again, the same measure for a shuffled and a shifted contact map; the correlation values in these cases are close to zero, and none are statistically significant.

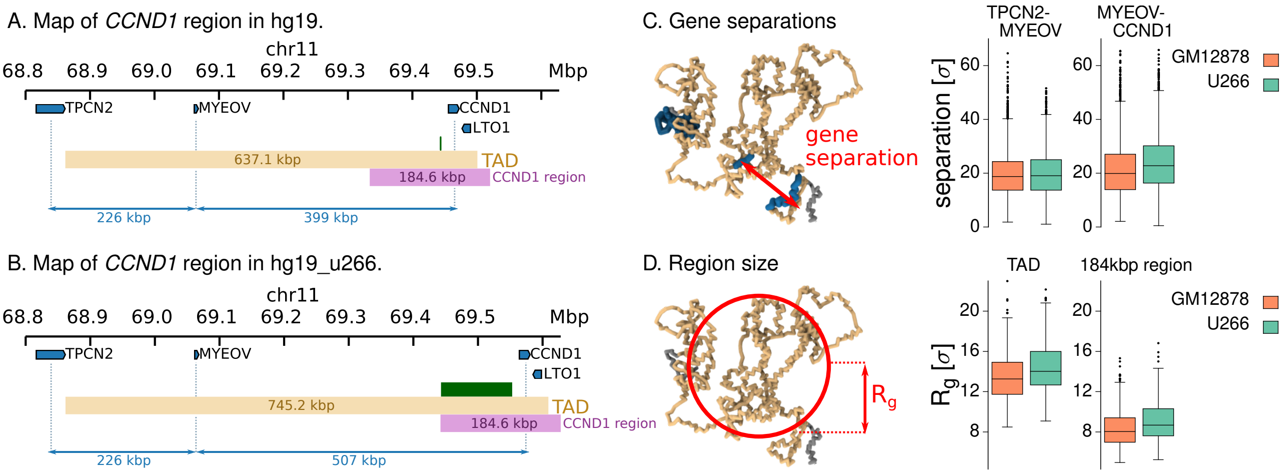

**Supplemental Figure S5. Single cell-like measurements from simulations.** In a given simulated structure we can make 3D distance measurements between specific points on the chromosome, *e.g.*, between the centres of mass of two gene bodies. We can also measure the 3D size of a given region. Each simulated structure represents a single cell, and we can obtain the distribution of these single cell measurements across the population of structures. **A.** Map showing gene locations within the hg19 genome, with their genomic separations indicated. The coloured blocks indicate the extent of the *CCND1*-TAD as detailed in the main text, and a 184.6 kbp region covering *CCND1*. **B.** A similar map shows the genes in the hg19_u266 genome which includes the insert (green block; the insert position is also shown in A). Note that due to the insert the genomic separation of *MYEOV* and *CCND1* increases, as does the linear size of the TAD. Another 184.6 kbp region covering *CCND1* is also shown (note it is the same size as in A). **C.** Separations of genes can be measured from each generated configuration. We define the position in 3D as the centre of mass of all beads within the gene. The box plot shows the distribution of separations across 4400 configurations for pairs of genes in the two cell lines (U266 with, and GM12878 without the insert). Only the *MYEOV*-*CCND1* separation shows a statistically significant difference ($p < 10^{-6}$ from a Kolmogorov-Smirnov two-sample test; the mean separation is 12% larger in U266 cells). Separations are given in simulation units $\sigma$, which represent approximately 21.8 nm (Buckle *et al.*, 2018). **D.** The 3D size of a given genomic region can be determined by its radius of gyration, $R_g$ (see Supplemental Methods for definition). Box plots show the distribution of $R_g$ measurements across 4400 configurations from each cell line. The left plot shows that the space taken up by the *CCND1*-TAD increases in U266 cells ($p < 10^{-5}$ from a Kolmogorov-Smirnov two-sample test; on average a 6% increase, note that the genomic length is also larger due to the insert). The right plot shows that a 184.6 kbp region covering *CCND1* in each cell line also increases in U266 cells (the genomic length is the same, but the chromatin states differ).

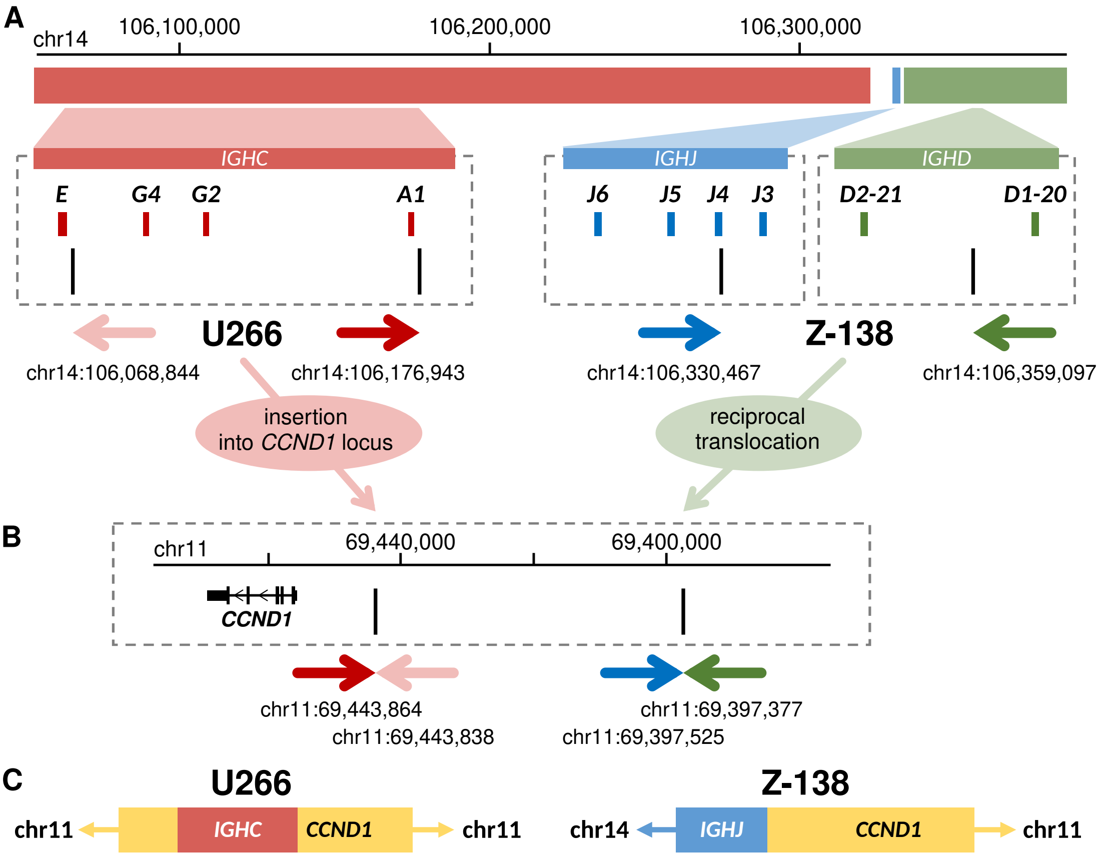

**Supplemental Figure S6. Schematic of the *IGH-CCND1* translocation in the U266 and Z-138 cell lines. A and B.** In the U266 cell line (left), two chromosomal break-points at *IGHC* (A, red) define a region that was cut and pasted into the *CCND1* locus (B). The *IGH* break-point positions suggest that this abnormality was likely formed as an error during class-switch recombination, resulting in insertion of *IGHC* next to *CCND1*. In the Z-138 cell line (right), exchange of DNA material between *IGH* and *CCND1* loci was confirmed. Two chromosomal break-points at the *IGH* locus were detected (A), in *IGHJ* (blue) and in *IGHD* (green). The break-point at chromosome 11 (B) showed that the part of this chromosome with *CCND1* becomes joined to the *IGHJ* end of chromosome 14. The architecture of this translocation suggests that it is a reciprocal translocation likely formed as an error during V(D)J recombination. Chromosomal break-points are shown as black vertical lines at each locus. Coloured horizontal arrows represent the translocation architecture; arrow colour indicates the parts of the chromosomes which are connected after the rearrangements (like colours are connected). **C.** The rearranged genome at *CCND1* is shown schematically for each cell line.

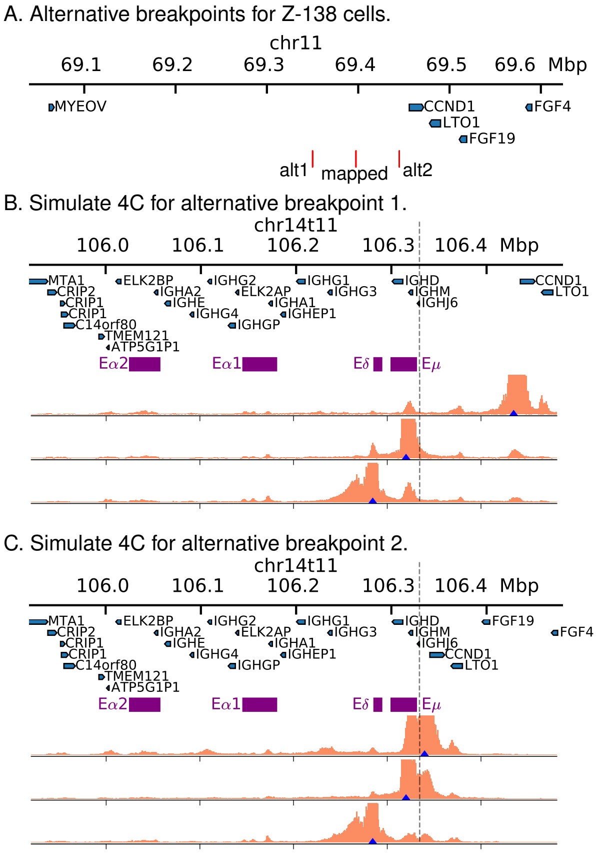

**Supplemental Figure S7. Simulating alternative break-points for Z-138 cells. A.** Map of chromosome 11 showing two alternative break-points and the mapped break-point. The first (alt1) is further upstream from the *CCND1* promoter than the mapped break-point, while the second (alt2) is closer to *CCND1* (*i.e.*, it is immediately upstream of the promoter). **B.** Simulated 4C data from three viewpoints, one at the *CCND1* promoter and two at DHS within the E$\delta$ and E$\mu$ super-enhancers. Simulations were performed for a rearranged genome with alternative break-point alt1, and input data from Z-138 cells. **C.** Similar plot but for the second alternative break-point. These results show that the frequency of interactions between *CCND1* and the enhancer increases the closer they are genomically. In the case of break-point alt1 (panel B), an additional DHS upstream of *CCND1* is brought from chromosome 11, which gives rise to a small additional peak of interactions.

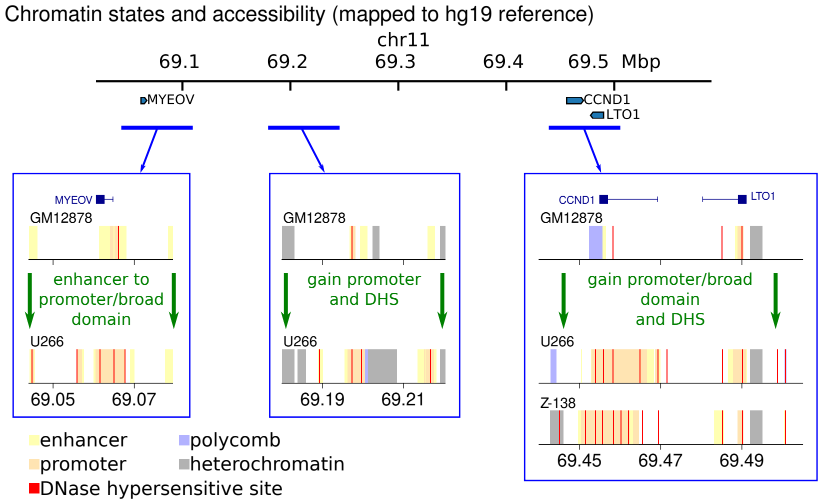

**Supplemental Figure S8. Differences between chromatin states at *CCND1* in the three cell lines.** Three chromosome 11 regions within the *CCND1*-TAD which show substantial chromatin remodelling in U266 and Z-138 cells are indicated with blue lines. Plots showing chromatin states as coloured blocks are shown for these regions as indicated by arrows; the data are mapped to the hg19 reference genome. Yellow, orange, blue and grey blocks indicate enhancer, promoter, polycomb and heterochromatin states respectively, with red lines indicating positions of DHS. Note that both enhancer and promoter states are associated with H3K27ac, and so are treated the same in simulations (open chromatin). Green text indicates how the chromatin is remodelled compared to GM12878.

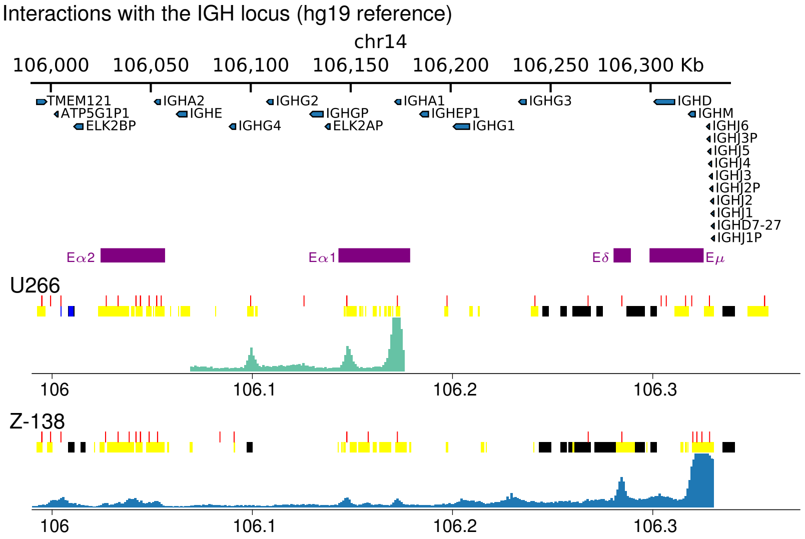

**Supplemental Figure S9. Interactions at the *IGH* enhancers in U266 *vs.* Z-138.** Plots showing 4C interactions across the *IGH* locus from a viewpoint at *CCND1* from simulations of U266 and Z-138 cells. These have been mapped back to the hg19 reference genome; only regions which are proximate to *CCND1* after the rearrangement show a signal. Enhancer positions are shown as purple blocks; red, yellow, blue and black blocks show positions of DHS, open chromatin, polycomb and heterochromatin regions. There are differences in the pattern of chromatin states and DHS between the two cell lines. Particularly, within E$\alpha$1, there are three DHS in Z-138, but only two in U266. In the latter these show strong interactions with *CCND1* due to their proximity to the gene. We also note that a DHS at around chr11:106,100,000 in U266 which interacts with *CCND1* is not present in Z-138; the chromatin state here is also different in the two cell lines. This suggests that the rearrangement not only leads to a remodelling of chromatin around *CCND1*, but also within the *IGH* enhancer region. It is important to note, however, that the aberrant rearrangements may arise from different errors in antibody production processes at different stages of B-cell differentiation (as discussed in Supplemental Methods section 5). From the interaction profiles we see that in U266, the two most frequently interacting DHS are at opposite ends of the E$\alpha$1 enhancer region. The DHS closest to the gene interacts most strongly. In Z-138, the strongest interaction with the *CCND1* promoter is with a cluster of DHS within E$\mu$ (*i.e.*, the closest enhancer genomically).

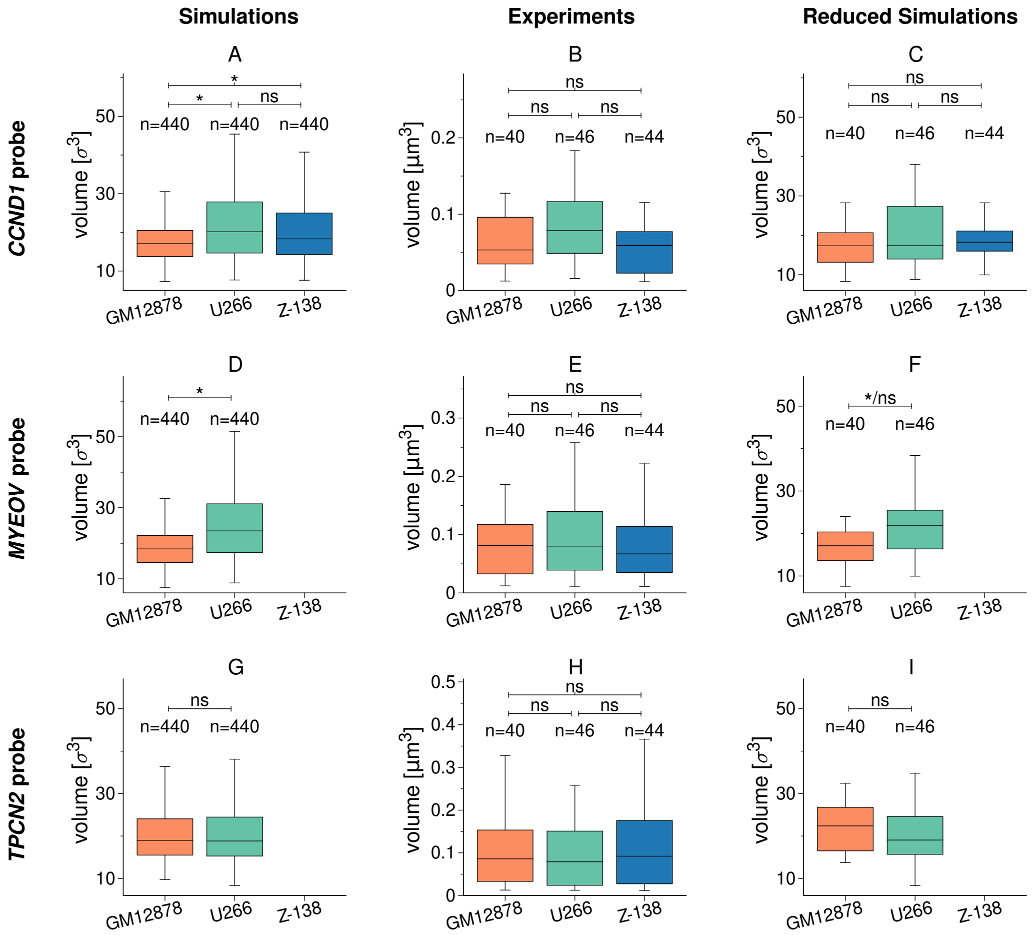

**Supplemental Figure S10. Fluorescence *in situ* hybridisation experiments.** Measurements of the volume of three BAC probes (genomic positions shown in Fig. 6D in the main text) were obtained for in GM12878, U266 and Z-138 cells (see Supplemental Methods for details). Distributions of volumes are shown as box plots, and the number of measurements $n$ is indicated above each box. A two-sample Kolmogorov-Smirnov test was used to determine the significance of the differences between the probe volumes in different cell line, as shown by brackets above the bars; '*' indicates that the null hypothesis could be rejected ($p < 0.01$), while 'ns' indicated that the null hypothesis could not be rejected and there is no significant difference between the distributions. **Left panels (A,D,G)** show simulated data (volumes obtained from the gyration tensor of the beads representing chromatin covered by the probe, see Supplemental Methods Section 4). **Middle panels (B,E,H)** show experimental data as in Fig. 6F in the main text. In our images, probe volume was determined by counting the contiguous voxels which have fluorescence intensity above a threshold; voxels had dimension 43×43×130 nm (volume $2.4\times10^{-4}$ $\mu$m$^3$). **Right panels (C,F,I)** show a reduced simulation data set. As detailed in Supplemental Methods Section 7, for better comparison between simulations and data, we chose a random subset of simulation measurements such that the total number of measurements matched the experimental data. In experiments we cannot differentiate between probes on rearranged and non-rearranged chromosomes, so to mimic this, in the U266 and Z-138 simulation cases, half of the included measurements were from GM12878 simulations. Box plots show one random subset of simulation measurements. When testing for significant differences, we repeated the test with 10 different random subsets for each case; '*/ns' indicates that for some random subsets there was a significant different, while for others there was not (suggesting that this number of measurements is on the cusp of being sufficient to differentiate the distributions). Each row of panels refers to a different probe: A-C for the *CCND1* probe; D-F for the *MYEOV* probe; and G-I for the *TPCN2* probe.

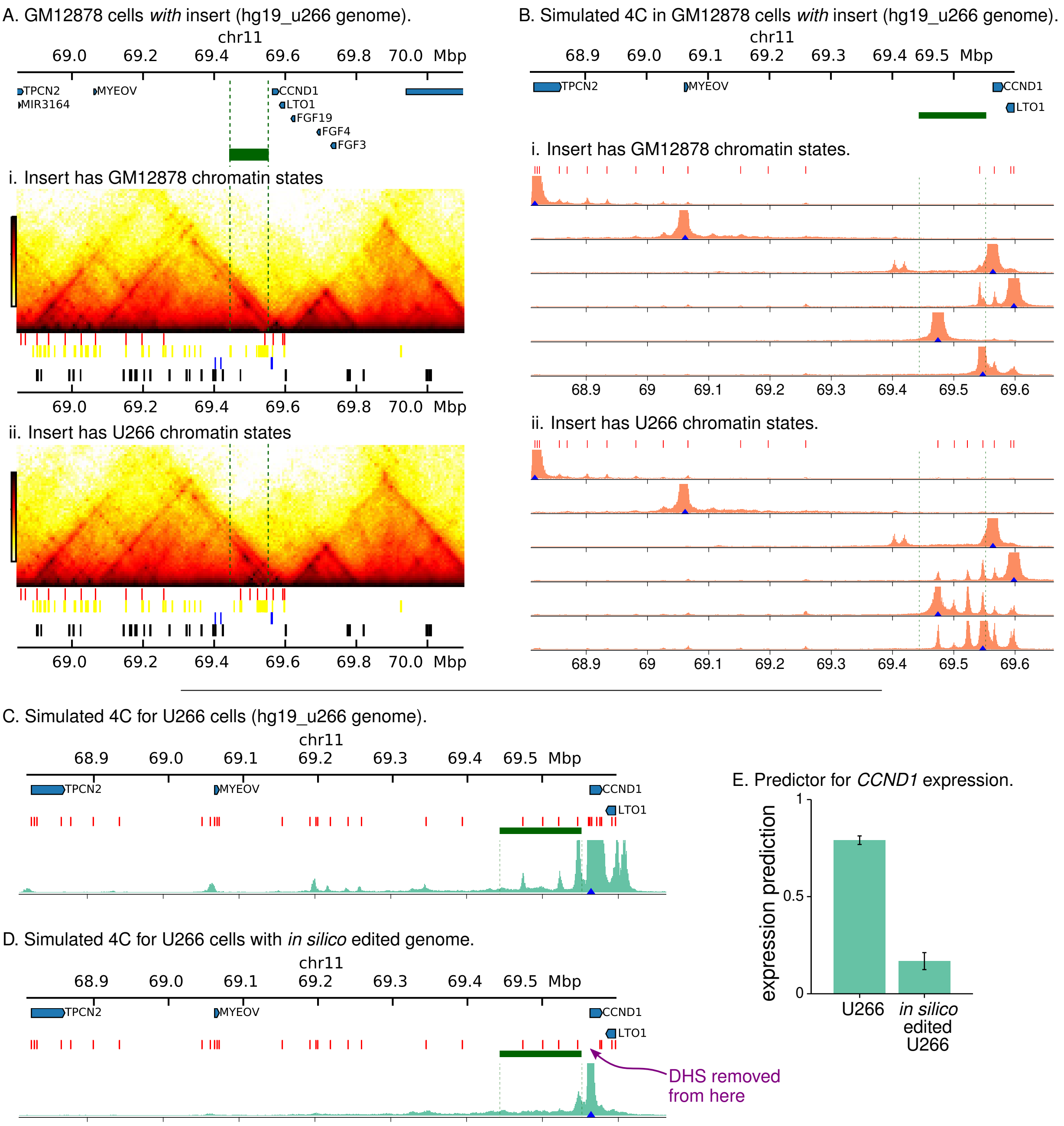

**Supplemental Figure S11. A and B: Simulating the *CCND1* locus in GM12878 cells, but adding the insert with U266 chromatin states. A.** Simulated Hi-C interaction maps for the *CCND1* locus with the *IGH* insert (hg19_u266 genome). Input data (chromatin states, DHS, and CTCF sites) for chromosome 11 are from GM12878 cells. The green block indicates the position of the insert, and coloured blocks show the input data as in Fig. 2 in the main text. In (i) the input data for the insert region are from GM12878 cells. In (ii) the input data for the insert region are from U266 cells; in this case the insert contains several DHS and a broad H3K27ac region, and so this constitutes a simulation where an active enhancer is inserted into the locus (but no further "epigenomic" rearrangement takes place). We note that there is very little difference between the maps; in (ii) there are some additional interactions within the insert region. **B.** Plots showing simulated 4C from the same simulations as in A. Positions of viewpoints are indicated by blue triangles, and red blocks above the plots show the positions of DHS. Vertical green dashed lines show the position of the insert. Interaction profiles from viewpoints at the promoters of *TPCN2*, *MYEOV* and *CCND1* do not show any difference between the two simulations. In the case of *TPCN2* this is because the gene is too far away genomically to interact with the insert; *MYEOV* and *CCND1* do not have DHS within their promoters, so there is nothing to drive interactions with the insert. The viewpoint at *LTO1* shows some additional interaction with the insert region in (ii): protein binding at the DHS within the *LTO1* promoter drives looping interaction with the DHS in the insert. In (ii) additional interactions between DHS within the insert are also observed. **C-E: Simulating the *CCND1* locus in U266, but editing the input data to remove DHS (active protein binding sites). C.** Plot showing simulation 4C for a probe at the *CCND1* promoter (blue triangle) for U266 cells, as in Fig. 3. in the main text. **D.** Similar plot but from a simulation where the input data has been edited to remove four DHS within the promoter and gene body of *CCND1* (as indicated by the purple arrow). A loss of interactions with nearby and more distant DHS is observed. **E.** Plot showing how the prediction for *CCND1* expression changes due to this "*in silico* epigenome editing".
